# Repeated evolution of a morphological novelty: a phylogenetic analysis of the inflated fruiting calyx in the Physalideae tribe (Solanaceae)

**DOI:** 10.1101/425991

**Authors:** Rocío Deanna, Maximilien D. Larter, Gloria E. Barboza, Stacey D. Smith

## Abstract

**PREMISE OF THE STUDY:** The evolution of novel fruit morphologies has been integral to the success of angiosperms. The inflated fruiting calyx, in which the balloon-like calyx swells to completely surround the fruit, has evolved repeatedly across angiosperms and is postulated to aid in protection and dispersal. Here we investigate the evolution of this trait in the tomatillos and their allies (Physalideae, Solanaceae), using a newly estimated phylogeny and a suite of comparative methods to infer evolutionary gains and losses.

**METHODS:** The Physalideae phylogeny was estimated using DNA sequences from four regions (ITS, LEAFY, *trnL*-*F*, *waxy*) using maximum likelihood and Bayesian Inference. Maximum likelihood model selection was used to determine the best fitting model of trait evolution. Using this model, we estimated ancestral states along with the numbers of gains and losses of fruiting calyx accrescence and inflation with Bayesian stochastic mapping. Also, phylogenetic signal in calyx morphology was examined with two metrics (parsimony score and Fritz and Purvis’ D).

**KEY RESULTS:** The well resolved phylogeny points to multiple taxa in need of revision, including the eight genera that are non-monophyletic as presently circumscribed. Model fitting indicated that calyx evolution has proceeded in stepwise fashion, from non-accrescent, to accrescent, to inflated. Moreover, these transitions appear to be largely irreversible. Among the 215 sampled Physalideae, we inferred 24 gains of fruiting calyx accrescence, 24 subsequent transitions to a fully inflated calyx and only two reversals. A median of 50 shifts were estimated in total across the clade from the ancestral non-accrescent calyx. Nonetheless, fruiting calyx accrescence and inflation show strong phylogenetic signal.

**CONCLUSIONS:** Our phylogeny greatly improves the resolution of Physalideae and highlights the need for taxonomic work. The analyses of trait evolution reveal that the inflated fruiting calyx has evolved many times and that the trajectory towards this phenotype is generally stepwise and directional. These results provide a strong foundation for studying the genetic and developmental mechanisms responsible for the repeated origins of this charismatic fruit trait.

---

Fruit evolution has long been considered a key contributor to the success of angiosperms, with bursts of morphological innovation closely tied to climatological events as well as the rise of frugivorous lineages of vertebrates (Tiffney, 1984; Eriksson et al., 2000; Knapp, 2002). Variation in fruit traits across taxa is often correlated with differences in dispersal mode (e.g., Gautier-Hion et al., 1985; Lomáscolo et al., 2010), which in turn, can lead to shifts in diversification rates (e.g., Beaulieu and Donoghue, 2013; Lagomarsino et al., 2016; Larson □ Johnson, 2016). Beyond their role in facilitating seed dispersal, fruits also serve to protect seeds from pathogens and predators (Tewksbury and Nabhan, 2001; Beckman and Muller-Landau, 2011) and promote successful germination (Traveset, 1998; Vander Wall, 2001).

From an evolutionary perspective, fruit morphology is known not only for its tremendous diversity but also the high degree of convergence. For instance, fleshy fruits have evolved repeatedly in a wide variety of angiosperm clades (e.g. Malphigiaceae, Davis et al., 2001; Rubiaceae, Bremer et al., 1995; Solanaceae, Knapp, 2002), often in relation to shifts in ecological niche (Bolmgren and Eriksson, 2005; Givnish et al., 2005). Even seemingly complex fruit traits, such as heteroarthrocarpy, have been gained and lost multiple times at recent phylogenetic scales (Hall et al., 2011; Marcussen and Meseguer, 2017). However, unlike with floral traits, such as symmetry and coloration (Preston and Hileman, 2009; Sobel and Streisfeld, 2013), the extent to which convergent transitions in fruit traits occur through similar genetic and developmental mechanisms remains little explored (Pabón-Mora et al., 2014; Ortiz-Ramírez et al., 2018; but see Avino et al., 2012).

Here we focus on a charismatic but understudied fruit trait, the inflated fruiting calyx, which has evolved repeatedly across angiosperms. Inflated calyces develop by accrescence after anthesis such that the fruit becomes completely enclosed upon maturation (He et al., 2004). This feature is found in at least 11 plant families, such as Malvaceae and Lamiaceae (Paton, 1990; Padmaja et al., 2014), although it is best known from the tomato family, Solanaceae, where it is referred to as a ‘chinese-lantern’ fruit or, more formally, the ‘inflated calyx syndrome’ (ICS; He et al., 2004; He and Saedler, 2005; Wang et al., 2015). This enlarged fruiting calyx has been proposed to aid in dispersal by acting as a tumbleweed (Knapp, 2002) or by providing flotation in flooded environments (Wilf et al., 2017). Pre-dispersal, the inflated calyx may also serve to protect the developing fruit from predators as well as from desiccation (Cedeño and Montenegro, 2004; Riss, 2009).

The evolution and development of inflated calyces has been studied in detail in only one clade, the tomatillos and their allies (tribe Physalideae, Solanaceae). Using comparative gene expression studies and transformation experiments, He and Saedler (2005) demonstrated that expression of a MADS-box transcription factor (*MPF2*) is required for the development of the dramatic inflated calyx in *Physalis*, and that overexpression of this gene in tomato can induce some degree of fruiting calyx accrescence. Subsequent studies across Physalideae revealed that many taxa which lack inflated calyces express *MPF2*, indicating that additional factors are required for development of the trait (Hu and Saedler, 2007). These and subsequent authors suggested that, given the shared expression of *MPF2* across Physalideae, ICS could be the ancestral state with multiple subsequent losses (Hu and Saedler, 2007; Zhang et al., 2012). Nonetheless, progress in reconstructing the history of gains and losses of this morphological innovation has been hampered by the sparse taxon sampling of Physalideae in existing phylogenies, which include only 37 % of the extant taxa (Särkinen et al., 2013).

In the present study, we aim to elucidate the evolutionary history of fruiting calyx inflation in Physalideae with a greatly expanded phylogeny and statistical comparative analyses of character transitions. This tribe contains the highest generic-level diversity in Solanaceae, with 29 genera and ca. 300 species arranged in three subtribes (Iochrominae, Physalidinae and Withaninae; Olmstead et al., 2008; Särkinen et al., 2013). Moreover, 13 of 19 Solanaceae genera with fruiting calyx inflation are placed in Physalideae, The wide variation in fruiting calyx form, from non-accrescent to greatly inflated (Fig. 1), has often been used for intergeneric delimitation (Hunziker, 2001; Sawyer, 2001; Li et al., 2013; Zamberlan et al., 2015), although phylogenetic studies suggest that these characters are homoplastic (Hu and Saedler, 2007). With a new phylogeny including 73 % of Physalideae species, we traced the evolution of the fruiting calyx accrescence and inflation to address the following questions: (i) is fruiting calyx inflation a convergent trait in Physalideae?; (ii) if so, how many times has this trait been gained or lost? (iii) can lineages move directly between non-accrescent and inflated states or do they tend to transition through intermediate stages of accrescence? The answers to these questions will provide insight into evolutionary accessibility of the lantern-like fruit form and lay the foundation for future studies at the genetic and development levels.

**FIGURE 1.**
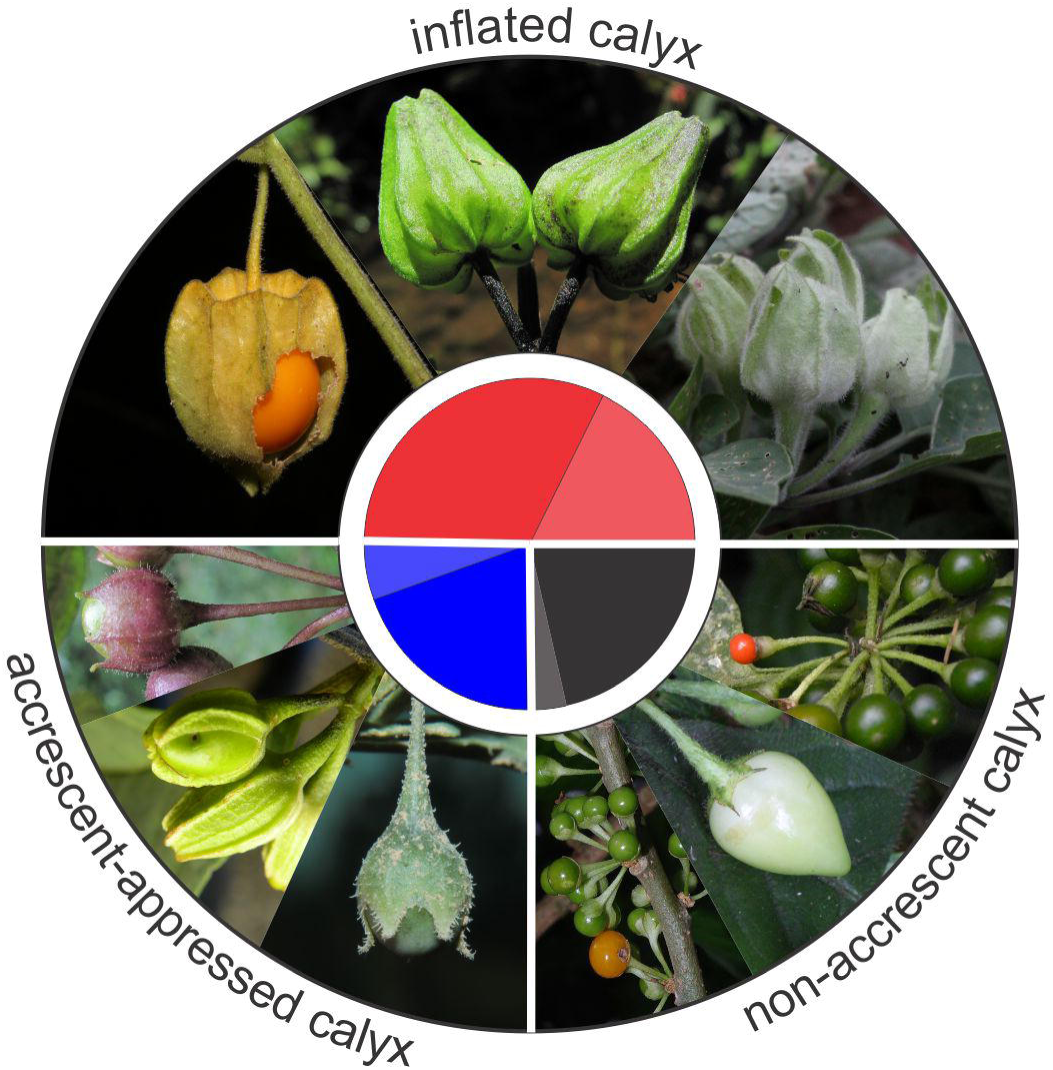
Distribution of fruiting calyx states across the tribe Physalideae. The size of the pie slices represents the proportion of taxa in each state, i.e. species with inflated calyces (red, 49.7%), non-accrescent calyces (black, 24.8%) and accrescent but still appressed calyces (blue, 25.5%). The darker shade in each pie slice corresponds to the percentage of taxa with that state sampled in the present study (64.4%, 86.3%, and 77.3%, respectively). Images from red to blue (moving clockwise) are *Physalis hederifolia* (Deanna *et al.* 209, photo by S. Carrasco), *Deprea pumila* (Orozco *et al.* 3890, photo by S. Leiva González), *Aureliana cuspidata* (Stehmann *et al.* 6457, photo by R. Deanna), *Witheringia solanacea* (Deanna 160, photo by R. Deanna), *Aureliana wettsteiniana* (Stehmann *et al.* 6448, photo by R. Deanna), *Iochroma arborescens* (Orejuela & Castillo 2697, photo by A. Orejuela), *Chamaesaracha coronopus* (Deanna *et al.* 237, photo by S. Carrasco), *Cuatresia exiguiflora* (Orozco *et al.* 3853, photo by G. E. Barboza), *Deprea sawyeriana* (Deanna & Leiva González 14, photo by S. Leiva González). Photos not to scale.

## MATERIALS AND METHODS

### Taxon sampling

–The ingroup sampling spanned 27 of the 29 genera of Physalideae and included 215 species of the 294 species plus 4 varieties (Appendix S1 and S2; see Supplemental Data with this article). The monotypic *Mellissia* Hook. f. and *Capsicophysalis* Averett & M. Martínez were the only genera not sampled. *Capsicum lycianthoides*, *Lycianthes inaequilatera*, and *Salpichroa tristis* (Appendix S1) were used as outgroups. Newly sampled plant material was either gathered from herbaria (CORD, CSU, MO, SI) or collected during several field trips to Argentina, Bolivia, Brazil, Colombia, Ecuador, Peru, and United States in the last ten years. Leaves were dried in silica and vouchers were prepared and housed at local herbaria of each country (Argentina: CORD; Bolivia: LPB; Brazil: BHCB; Colombia: COL, JBB, PSO; Ecuador: LOJA, QCA, QCNE, QUSF; Peru: HAO, HUT; United States: COLO, CSU, MO). We also obtained already extracted DNA from L. Bohs, R. Olmstead, and L. Freitas.

### Phylogenetic reconstruction of Physalideae

–We used de novo (407, ca. 55 %) and published (339, ca. 45 %) sequences from four regions to estimate relationships within Physalideae (Appendix S1, including GenBank accession numbers): the nuclear regions internal transcriber spacer (ITS), granule-bound starch synthase (GBSSI or *waxy*) gene, the second intron of LEAFY (LFY), and the chloroplast spacer *trnL*-*F.* GBSSI regions previously sequenced by Whitson and Manos (2005) were not included in the analyses because they only comprised from exon 8 to 10, whereas we are using from exon 2 to 9 for most taxa. Taxa coverage was 92.8 % for ITS, 77.9 % for LFY, 78.4 % for *waxy*, and 87 % for the chloroplast fragment (Appendix S3). DNA extractions were done following a modified 2 x CTAB procedure (Doyle and Doyle, 1987); primers and PCR conditions followed previous work (Smith and Baum, 2006; Deanna et al., 2018).

Sequence quality was inspected using GENEIOUS v4.6.1 (Drummond, et al., 2006), and sequence alignments were performed in MEGA 6 (Tamura et al., 2013) using the MUSCLE algorithm (Edgar, 2004) followed by manual adjustments. For *trnL*-*F*, a variable repeat region towards the 5’ end of the intergenic spacer was removed because this is where putative pseudogenic copies of *trnF* have been found in *Solanum* (Poczai and Hyvönen, 2011). Gene trees were estimated individually for each region with maximum likelihood (ML) in RAxML v.8 (Stamatakis, 2014) on the CIPRES server (Miller et al., 2010). We implemented the GTR + GAMMA model and used the rapid bootstrap (BS) algorithm with 1000 replicates to assess nodal support. Trees were compared across genes to identify areas of hard incongruence (BS > 70%; Mason-Gamer and Kellogg, 1996).

Given the absence of hard incongruence, we conducted ML and Bayesian analyses on the combined dataset. Matrices were concatenated with SequenceMatrix 1.8 (Vaidya et al., 2011) and partioned by gene before analysis. We also identified unstable tips based on the ML bootstrap analyses using the software RogueNaRok (Aberer et al., 2013). Two iterations of RogueNaRok were run with settings according to Särkinen et al. (2013), and rogue taxa were removed after each iteration, resulting in the pruning of 10 tips in total. We also excluded the voucher R. Deanna 143 (which morphologically matches to the original description of *Cuatresia harlingiana* Hunz.) given its phylogenetic position outside of Physalideae. However, we included sequences of a voucher previously identified as *C. harlingiana* (Smith and Baum, 2006; Deanna et al., 2017, 2018), which does fall within *Cuatresia* and appears to belong to an undescribed taxon (appearing here as *Cuatresia* sp.).

The final combined matrix included 7988 bp of aligned sequences of 222 taxa, including outgroups. We performed ML phylogenetic inference partitioned by gene using RAxML according to the parameters used for individual region analyses (see above) on the CIPRES server (Miller et al., 2010). Bayesian analyses were conducted for the combined dataset with four partitions in BEAST 2 (Bouckaert et al., 2014), also on the CIPRES server. Best models of substitution were incorporated for each partition according to a previous selection with the Akaike Information Criterion (AIC) using jModelTest 2.1.3 (Appendix S3; Posada and Crandall, 1998; Darriba et al., 2012). Two independent BEAST analyses were run for fifty million generations each with tree sampling every 1000 generations, using an uncorrelated lognormal relaxed clock model to describe the branch-specific substitution rates (Drummond, et al., 2006). We used a Birth-Death tree prior, which accounts for both speciation and extinction (Gernhard, 2008), and a constraint of monophyly for all species excluding *Salpichroa tristis.* Convergence and stationarity of the parameters were inspected using Tracer v1.7 (Rambaut et al., 2018), targeting minimum effective sample sizes (ESS) of at least 200. The initial 20 % of trees were discarded as burn-in, and the results were combined using LogCombiner as implemented in the BEAST package. The phylogenetic relationships were summarized in a maximum clade credibility (MCC) tree, and their posterior probabilities (PP) for all nodes were derived using TreeAnotator v2.4.7. The trees were visualized in FigTree v.1.4.3 (Rambaut, 2016).

### Codification of fruiting calyces

–All fruiting calyces from taxa included in the phylogeny were scored using specimens housed at herbaria (COL, COLO, CORD, CSU, MO, SI), the JSTOR Plants database, and the literature (Appendix S2). Following Hu and Saedler (2007), we scored a fruiting calyx as accrescent-appressed when there is an increase in calyx length of 50 % or more from flower to fruit stage (e.g. *Brachistus stramoniifolius*), or the berry is entirely covered but there is not a space between calyx and berry (e.g. *Cuatresia exiguiflora*). Fruiting calyx was coded as non-accrescent when it grows less than 50 % from flower to fruit stage (e.g. *Witheringia solanacea*), and as inflated when the fruit is entirely enclosed by the calyx and there is also a space between calyx and berry (e.g. *Physalis peruviana*; see matrix in Appendix S4). Note that following this definition, species of *Iochroma* are coded as non-accrescent despite being described as often having accrescent calyces (Hunziker, 2001; Smith and Baum, 2006; Lezama Escobedo et al., 2007; Cueva Manchego et al., 2015). In *Iochroma*, accrescence is usually less than the 50 % of the length present at the flowering stage. In a handful of species (e.g. *I. calycinum*, *I. barbozae*; Khan et al., 2012a; Leiva González et al., 2013), the fruiting calyx covers the berry (or nearly so), but this is due to the large size of the flowering calyx.

### Testing for phylogenetic signal

–We implemented two metrics to examine the level for phylogenetic signal in fruiting calyx morphology. First, we calculated the parsimony score using the *parsimony* function in the {phangorn} R package (Schliep, 2011). Second, we computed Fritz and Purvis’ D (FPD, Fritz and Purvis, 2010), a metric which captures the sum of sister clade differences, also available in {phangorn}. The FPD statistic takes a value of 1 if the trait has a phylogenetically random distribution and 0 if the trait has evolved under Brownian motion (Fritz and Purvis, 2010). For both measures, we tested whether the observed values differed those expected by chance (no phylogenetic signal) as well as those expected under Brownian motion. In the former case, the null distribution was created by randomly reshuffling the tip states 1000 times, and in the latter case, by evolving these traits on the phylogeny under a Brownian motion model 1000 times. These null distributions were created with the *treestat* function in the {phylometrics} package (Hua and Bromham, 2016). Traits with phylogenetic signal are predicted to differ significantly from the random distribution (p<0.05) but not the distribution expected under Brownian motion. As the FPD statistic can only be applied to binary traits, we considered fruit accrescence and inflation separately (Appendices S5 and S6), while for parsimony, we were able to examine them jointly as three-state character (Fig. 1). These analyses were conducted using the MCC tree.

### Reconstructing the evolutionary transitions to fruiting inflated calyces

–We estimated the history of fruit calyx evolution across Physalideae using maximum likelihood and Bayesian approaches. We first compared the fit of alternative models of trait evolution using the {ape} package in R (Paradis et al., 2004) and the MCC tree from the BEAST analyses. We considered six models with the first having transition rates between all states free to vary (the all rates different model) and the second with all rates equal. We then fit four stepwise models, where lineages move from non-accrescent to inflated through the intermediate state of accrescent-appressed. Model 3 has all steps being reversible while the last three models have one or more of these steps constrained to be irreversible (Table 1). Model selection was conducted with the Akaike Information Criterion (AIC) score, with the best model having a score at least two AIC units lower than the model with the next lowest AIC score (Burnham and Anderson, 2002).

**Table 1.** Comparison of likelihood models tested for fruiting calyx accrescence and inflation, including log-likelihood (lnLik) and Akaike Information Criterion (AIC) scores. The lowest AIC score is bolded. The character states are: 0 = non-accrescent, 1 = accrescent-appressed, 2 = inflated fruiting calyx, and thus q_01_, for example, denotes the transition rates from non-accrescent to accrescent-appressed.

| Model tested | Constraints | Free parameters | lnLik | AIC |
| --- | --- | --- | --- | --- |
| 1. All rates different | -- | 6: $q_{01}$, $q_{10}$, $q_{02}$, $q_{20}$, $q_{12}$, $q_{21}$ | -109.364 | 230.727 |
| 2. Equal rates | $q_{01} = q_{10} = q_{02} = q_{20} = q_{12} = q_{21}$ | 1: $q$ | -123.646 | 249.291 |
| 3. Stepwise reversible | $q_{02}=0$, $q_{20}=0$ | 4: $q_{01}$, $q_{10}$, $q_{12}$, $q_{21}$ | -109.374 | 226.747 |
| 4. Stepwise 0-1 irreversible | $q_{02}=0$, $q_{20}=0$, $q_{10}=0$ | 3: $q_{01}$, $q_{12}$, $q_{21}$ | -109.374 | **224.747** |
| 5. Stepwise 1-2 irreversible | $q_{02}=0$, $q_{20}=0$, $q_{21}=0$ | 3: $q_{01}$, $q_{12}$, $q_{10}$ | -111.630 | 229.259 |
| 6. Stepwise irreversible | $q_{02}=0$, $q_{20}=0$, $q_{21}=0$, $q_{10}=0$ | 2: $q_{01}$, $q_{12}$ | -111.630 | 227.259 |

Using the best fitting model, we next estimated ancestral states and the number of transitions between states with Bayesian stochastic mapping (SM). Through rounds of simulation (‘realizations’), SM generates a sample of histories of discrete character evolution on a phylogeny that should approximate the posterior distribution of histories (Huelsenbeck et al., 2003). In order to incorporate phylogenetic uncertainty, we performed 500 simulations of character history on a sample of 100 trees from the BEAST analysis with the combined dataset. The simulations, carried out with the *make.simmap* function in {phytools} package (Revell, 2012), were summarized on the MCC tree to provide the posterior probability of each state at each node. We also estimated the median number of changes for each transition type from the histories and computed 95% credibility intervals using the *hdr* function from the {diversitree} package in R (FitzJohn, 2012).

## Results

### Phylogeny of Physalideae

–Our final combined matrix had a taxon coverage of 0.84 % (Appendix S3) and comprised 215 species of Physalideae. This represents 73.1 % of the total species within the tribe and 55 % of the species within *Physalis.* The plastid *trnL*-*F* and the nuclear region ITS were the most densely sampled, whereas ITS contributed most parsimony-informative characters (Appendix S3). Hard incongruence was not found among gene trees (Appendix S7). The maximum likelihood and Bayesian topologies were largely congruent (Fig. 2 and Appendix S8, respectively) and showed strong to moderate support for Physalidinae (BS = 63 %, PP = 1) and Iochrominae (BS = 100 %, PP = 1), which is resolved as sister to the remaining Physalideae taxa (BS = 89 %, PP = 1). The previously proposed subtribe Withaninae (Olmstead et al., 2008; Särkinen et al., 2013) does not appears to be monophyletic but instead divided amongst two clades with the Hawaiian *Nothocestrum* and allied Old World genera more closely related to Physalidinae than other members of Withaninae. Moreover, eight of the 27 sampled genera are non-monophyletic as presently circumscribed (e.g. *Iochroma*, *Cuatresia*, *Physalis*).

**FIGURE 2.**
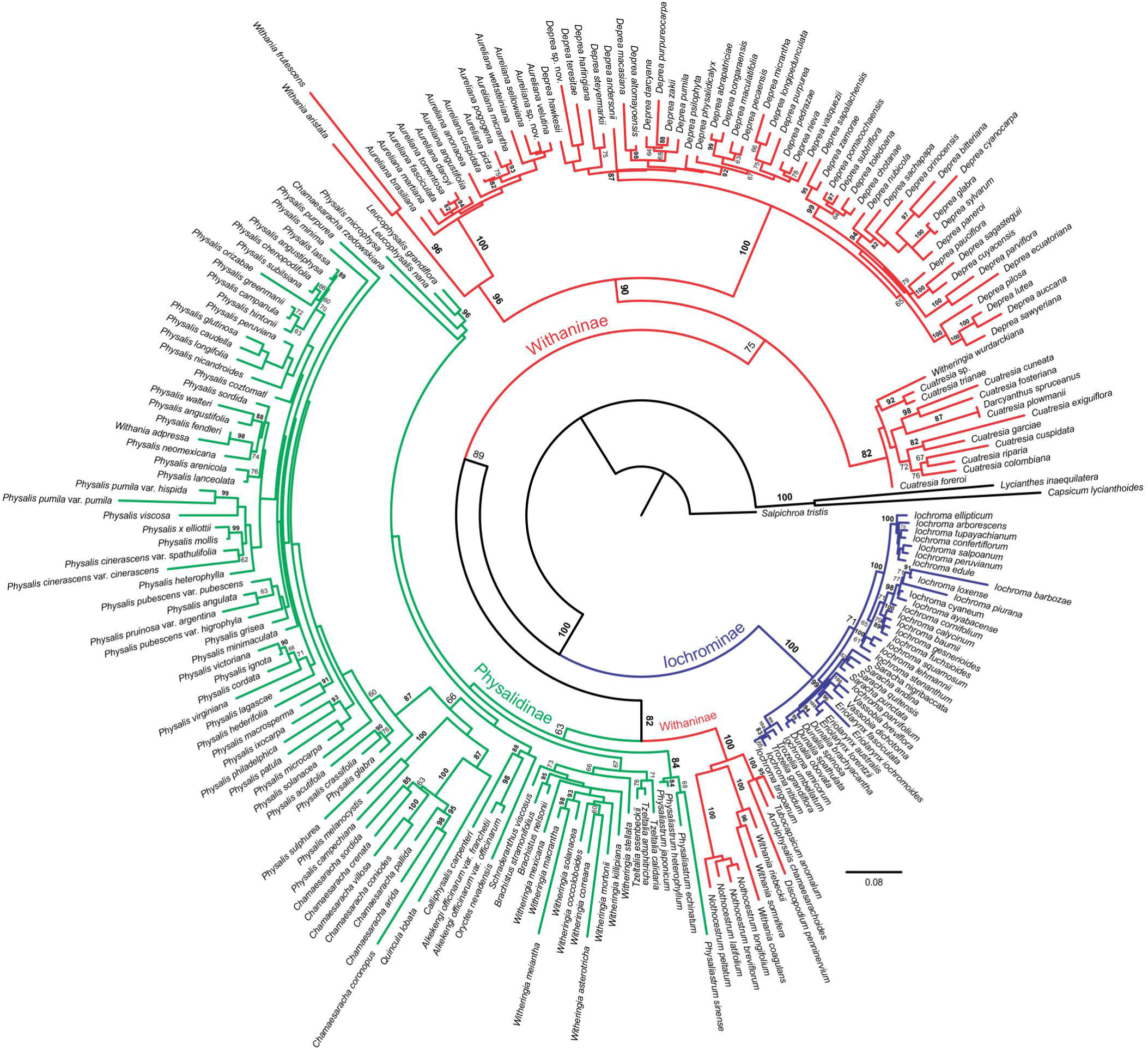
Phylogenetic relationships of Physalideae based on a maximum likelihood analysis of the combined dataset of four markers (ITS, LFY, *trnL*-*F*, and *waxy*). Bootstrap support (BS) values > 60 % are given above each branch, and bold numbers indicate BS > 80 %. Differentially coloured branches correspond to the subtribes proposed by Olmstead et al. (2008) and followed by Särkinen et al. (2013).

### Testing for phylogenetic signal of accrescent and inflated fruiting calyces

–We found strong phylogenetic signal for fruiting calyx accrescence and inflation with both implemented approaches. These traits have a significantly lower parsimony score and lower FPD compared to the random null distribution, suggesting that species with accrescent and inflated calyces are more closely related than expected by chance (Table 2). Consistent with this result, neither of the traits significantly differed from expectations under Brownian motion of evolution along the phylogeny (Table 2).

**Table 2.** Phylogenetic signal metrics calculated on Physalideae species with accrescent and/or inflated calyx. Bolded values indicate the statistics that were significantly lower than the random distribution of traits or significantly greater than Brownian motion evolution (p < 0.05). ^∗^FPD can adopt negative values up to −0.5 when the phylogenetic signal is high (Fritz and Purvis, 2010).

| Trait | Parsimony score (PS) | P-value of observed vs. random distribution | P-value of observed vs. Brownian motion evolution | Fritz and Purvis' D (FPD) | P-value of observed vs. random distribution | P-value of observed vs. Brownian motion evolution |
| --- | --- | --- | --- | --- | --- | --- |
| Fruiting calyx accrescence and inflation | **33** | **0.000** | 0.910 | NA | NA | NA |
| Fruiting calyx accrescence | **14** | **0.000** | 0.960 | **-0.483\*** | **0.002** | 0.961 |
| Fruiting calyx inflation | **20** | **0.003** | 0.934 | **-0.356\*** | **0.000** | 0.949 |

### Evolutionary transitions to fruiting inflated calyces

–The best fitting maximum likelihood model for fruiting calyx evolution was the stepwise model with transitions between accrescent and non-accrescent fruiting calyces being irreversible (reverse transition rate not different from zero). This model had the lowest AIC score and was greater than two AIC units lower than any competing model (Table 1; Appendix S9). Our stochastic mapping simulations with this model estimated a median of 50 changes across the clade (95% HDR = 44.56–56.04). Among these changes, shifts from non-accrescent to accrescent-appressed calyces and accrescent-appressed to inflated calyces were inferred to occur at roughly equal frequencies (median = 24 (19.94–29.09) vs. 24 (19.96–27.71), Appendix S10). Loss of inflation to an accrescent-appressed calyx was infrequent (median = 2, 95% HDR = 0–3.93; Appendix S10). The ancestral state of the tribe was estimated by SM as non-accrescent in all stochastic maps (100% posterior probability, Fig. 3). Similarly, high support was inferred for this ancestral state at many nodes throughout the phylogeny, revealing multiple independent gains of accrescence and inflation (Fig. 3).

**FIGURE 3.**
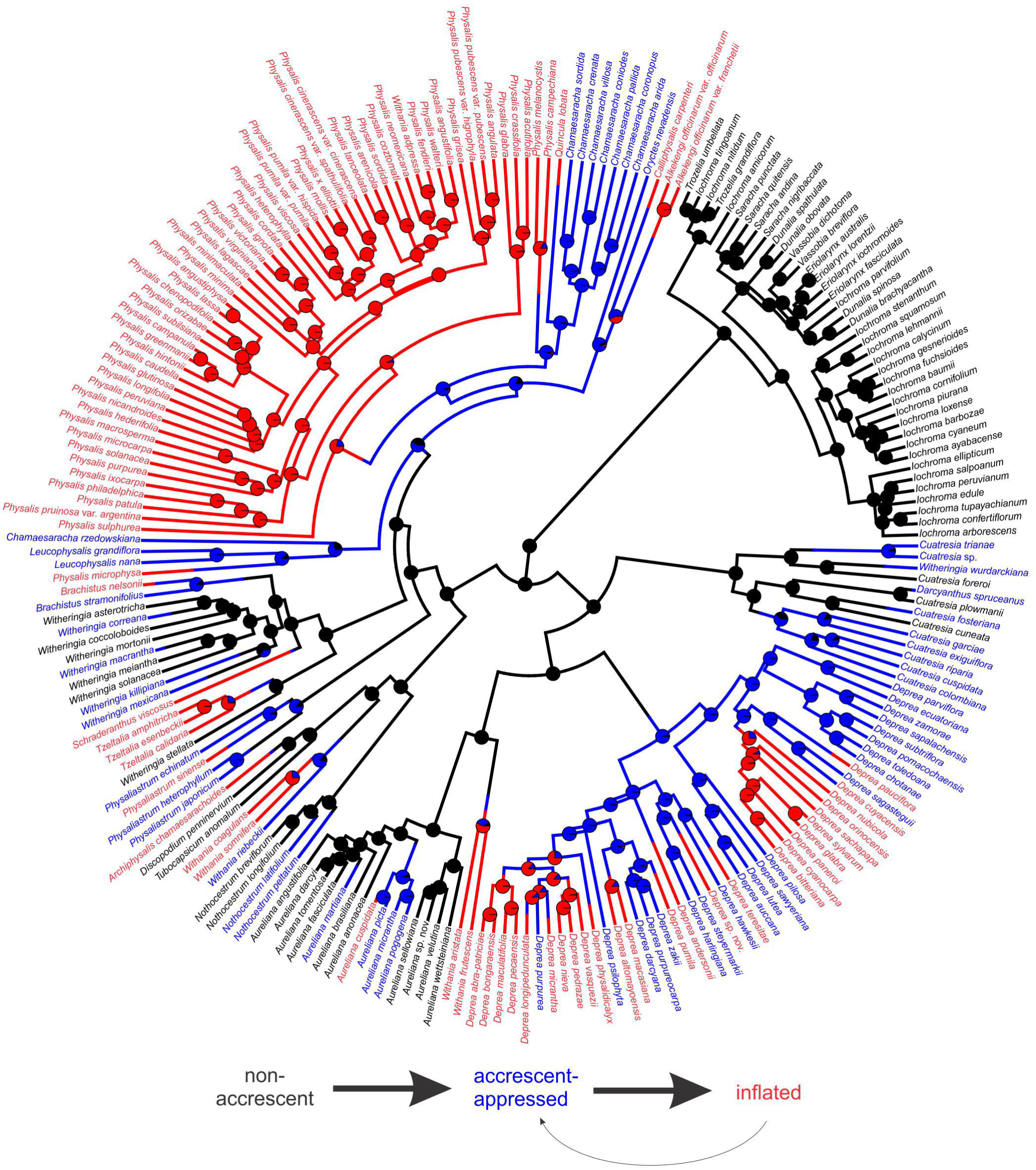
Reconstruction of fruiting calyx evolution in the Physalideae tribe. Topology is provided from four-gene BEAST analyses of 219 taxa. Circles at nodes indicate the posterior probabilities from stochastic mapping and tip label colors represent tip states, with red, blue and black representing inflated, accrescent-appressed, and non-accrescent fruiting calyces, respectively. On the bottom, transitions between states are represented with arrows proportional to number of estimated changes (see Appendix S10).

## DISCUSSION

### Phylogenetic relationships and taxonomy of Physalideae

–We present the first well-resolved and densely sampled phylogeny of the Physalideae tribe. This data set is a significant expansion compared with previous studies (e.g. 33 species of *Physalis* in Zamora-Tavares et al., 2016, vs 53 taxa here), and the sampling covers most of the taxonomic, morphological, and geographic variation within this group. Although some parts of the tree (e.g. within *Physalis*) will require additional data for better resolution, our results have recovered many previously proposed relationships as well as several new ones, which we briefly review here.

Starting with the monophyletic and well-studied Iochrominae, only three out of six genera are monophyletic, even after recent nomenclatural changes (Shaw, 2018a; b). The crossability among genera, high convergence in traits used to delimitate generic taxonomy, and the comparative lack of karyological variation (Smith and Baum, 2006; Smith et al., 2008; Shaw, 2018b) suggest that combining the genera into a single monophyletic *Iochroma* may be the most stable solution. During the last 20 years, 19 new species of *Iochroma* and one *Saracha* have been described (e.g. Leiva Gonzalez et al., 2003; Leiva González and Lezama, 2005; Lezama Escobedo et al., 2007; Fernandez-Hilario and Smith, 2017) but no key for the entire group has been proposed, increasing the necessity of a full taxonomic revision.

The subtribe Withaninae also presents taxonomic challenges, both at the subtribal and generic levels. This subtribe was originally circumscribed by Olmstead et al. (2008) to contain seven small genera, which were all Old World except for the South American *Aureliana.* Our analysis provides strong support for the non-monophyly of the type genus, *Withania*, with three species (*W. coagulans*, *W. riebeckii*, and *W. somnifera*) closely related to other taxa in Withaninae sensu Olmstead et al. (2008) and the other two species (*W. aristata* and the type species *W. frutescens*) closely related to *Aureliana.* In a prescient review, Hepper (1991) pointed out that these two western African species, *W. aristata* and *W. frutescens*, are morphologically unlike others in *Withania* and suggested that their closest relatives may instead be across the Atlantic. Beyond the rearrangement of Withaninae necessitated by this apparent split within *Withania*, most of the genera which have been placed in the subtribe are monophyletic (or nearly so) given extensive taxonomic work in recent years (Zamberlan et al., 2015; Deanna et al., 2018).

The largest subtribe Physalidinae, with 12 genera, was recovered as a monophyletic group although relationships among and within the genera are complex and, in some cases, unresolved. One complicating factor is the large number of monotypic genera (*Alkekengi*, *Calliphysalis*, *Oryctes*, *Quincula*, *Schaderanthus*), some of which are nested within other genera. Nonetheless, several of the affinities that we uncovered have been proposed by previous authors using morphological data (e.g. between *Brachistus* and *Witheringia* (Hunziker, 1969; between xerophytic *Chamaesaracha* but excluding *C. rzedowskiana*, Averett, 1973; Turner, 2015), suggesting viable avenues for future taxonomic rearrangements. Perhaps the greatest challenge will be estimating relationships within *Physalis*, which remain largely unclear in this study as they have in previous (Whitson and Manos, 2005; Zamora-Tavares et al., 2016). The lack of resolution within this clade may reflect a history of rapid diversification and hybridization, which will likely be elucidated only with phylogenomic approaches (e.g. Stenz et al., 2015).

### Repeated evolution of fruiting calyx accrescence and inflation

–Our analyses demonstrate that the highly-inflated fruiting calyx considered so characteristic of *Physalis* has evolved repeatedly in Physalideae. While previous studies had suggested homoplasious patterns in fruiting calyx variation in the tribe (Whitson and Manos, 2005; Hu and Saedler, 2007), we provide the first estimates of the numbers of gains and losses, with ca. 24 gains of accrescence, 24 subsequent gains of inflation and 2 reversals from inflation to the accrescent-appressed state (Fig. 3). Despite these many gains of calyx accrescence and inflation, we recovered significant phylogenetic signal in these traits overall. Indeed, the character states appear clustered on the phylogeny, with some large clades (e.g. Iochrominae, *Physalis*) being invariant in the degree of fruiting calyx accrescence.

The many independent origins of calyx inflation may have occurred through recurring modifications of the shared underlying pathway, which is well studied in several Physalideae. The development of ICS requires the expression of *MPF2*-like MADS-box transcription factors in flowering calyces (He and Saedler, 2005). Nonetheless, *MPF2* expression in the calyx is widespread across taxa with and without ICS in Physalideae and even in Capsiceae (Hu and Saedler, 2007), suggesting the development of ICS is determined by other factors. Indeed, the effect of *MPF2* on calyx morphology appears to hinge on interactions with cytokinin and gibberellin, which are released upon fertilization (He and Saedler, 2007; Khan et al., 2012b). Thus, genetic changes which modify these hormonal signals, *MPF2* expression, or *MPF2* function could all contribute to variation in calyx inflation (Riss, 2009). Comparative molecular and developmental studies to-date implicate both regulatory and structural mutations in *MPF2*-like genes (Hu and Saedler, 2007; Riss, 2009; Khan et al., 2009), coupled with shifts in copy number due to the many ploidy changes in the tribe (Iqbal and Datta, 2007; Deanna et al., accepted).

Inflated calyces have convergently evolved in many taxa outside of Solanaceae, although the possibility that these rely on the same genetic pathway has not been explored. The 11 families with highly accrescent calyces, in which the fruits may be berries, drupes or capsules, are spread across eudicots, from rosids (e.g., Caryophyllaceae, Malvaceae, Aptandraceae) to asterids (e.g., Lamiaceae, Boraginaceae, Campanulaceae) (Paton, 1990; Francis, 2000; Gottschling and Miller, 2006; Wilf et al., 2017). Solanaceae is the only family in which the developmental genetics of the trait has been studied in detail (Wang et al., 2015). Intriguingly however, overexpression of *MPF2*-like genes in *Arabidopsis* results in enlarged and persistent calyces (Khan et al., 2013) and the *MPF2*-like promoters from Physalideae are able to drive sepal-specific gene expression, also in *Arabidopsis* (Khan, et al. 2012b). These patterns suggest that many elements of networks regulating sepal growth are widely conserved, raising the possibilty that the evolution of inflated calyces in other clades has involved similar mechanisms.

### Loss vs. gain of inflation through a stepwise model of evolution

–Our comparative analyses indicated that evolution of the inflated calyx proceeds in directional fashion, starting from the non-accrescent state, moving first to an accrescent but appressed state before finally becoming inflated. This pattern contradicts the hypothesis that, given the complex developmental pathway required to produce ICS, inflation should be easier to lose than to gain (Hu and Saedler, 2007). This frequent and directional transitions toward inflation suggest not only that the trait is genetically accessible (perhaps given the background of *MPF2* expression in Physalideae calyces) but also that inflation is generally retained by lineages in which it evolves. Still, the adaptive advantages which could favor the fixation of this trait (e.g. protection from desiccation, deterrence of predators, enhanced dispersal) have been largely untested (but see Wilf et al., 2017). In fact, the only evidence for adaptive evolution of ICS comes indirectly from molecular studies, which have estimated positive selection acting on *MPF2*-like genes in *Withania* and *Physalis* (Khan et al., 2009; Zhang et al., 2012).

The retention of ICS following its evolution may reflect not only selective advantages, but also developmental constraints acting on reversals. Ablation experiments in two ICS taxa (*Physalis* and *Withania*) reveal a complex crosstalk between the calyx and fruit development at early stages, wherein removal of sepals prior to fertilization completely abolish fruit setting (He and Saedler, 2007; Khan, et al. 2012b); even ablations at later stages result in the development of smaller berries. These results suggest that genetic changes which reduce sepal size in ICS taxa might also reduce fruit size, which would presumably carry negative consequences for plant fitness. In the future, it would be valuable to conduct similar ablation experiments across Physalidae with non-accrescent, accrescent-appressed, and inflated calyxes to determine whether the negative effect of calyx damage on fruit development scales with the degree of accrescence of the fruiting calyx.

Despite the strong directionality inferred from our analyses, it is important to note that such patterns may be confounded by state-dependent differential diversification. For example, the abundance of inflated calyces (as in Physalideae) can occur through biased transitions toward this state or by increased diversification of lineages with the state (Ng and Smith, 2014). A thorough analysis of the effects of calyx evolution on speciation and extinction rates will require a larger phylogeny (Beaulieu and O’Meara, 2016), ideally at the family level and including all of the remaining genera (6) and species (76) with ICS. Diversification analyses would also benefit from new divergence time estimates in light of the recent discovery of Eocene lantern fruit fossils. These fossils, placed in crown group *Physalis*, are dated to 52.2 mya, which is roughly the age inferred for the entire crown group Solanaceae in previous work (Särkinen et al., 2013; De-Silva et al., 2017). This contrast highlights the need for a complete reassessment of Solanaceae fossils (Wilf et al., 2017; Särkinen et al., 2013, 2018), together with a new family-wide dating analysis including all reliable fossil taxa.

## CONCLUSIONS

Our phylogeny provides a starting point for re-circumscription of taxa and lays the foundation for ongoing research into morphological diversification of Physalideae and its spread around the globe. The charismatic lantern fruits, characteristic of the genus *Physalis*, have evolved repeatedly among its closely relatives in Physalideae. In each case, lineages have moved stepwise towards the inflated calyx, with many extant lineages exhibiting intermediate states of accrescence. This well resolved evolutionary history for Physalideae, together with the growing knowledge of fruit developmental pathways, will facilitate future work to trace the genetic changes that lead to ICS and may also explain the apparent directionality of transitions toward this morphological novelty.

## ACKNOWLEDGEMENTS

The authors thanks L. Bohs, L. Freitas and R. D. Olmstead for DNA samples, J. Stone, C. Carrizo García, Z. Zhang and F. Chiarini for leaf samples, herbaria staff from COLO, CORD, FLAS, MO, and SI for permission to extract leaf samples from specimens. We also greatly appreciate the help of C. Pretz, S. Carrasco and L. Fernández during fieldwork, and insightful suggestions of J. Ng on R scripts. Photos were kindly provided by A. Orejuela, S. Carrasco and S. Leiva González, and S. Liu was very helpful with translating Chinese for Asian taxa. The authors gratefully acknowledge support from the National Science Foundation (grants #1553114 to S.D.S., and #1556666 to P. Wilf), Consejo Nacional de Investigaciones Científicas y Técnicas (CONICET, grant PIP 00147), Agencia Nacional de Promoción Científica y Tecnológica (FONCyT, grant PICT 2017-2370) and SECyT (Universidad Nacional de Córdoba, Argentina). The first author also thanks to Fulbright and Ministerio de Educacion y Deportes (Argentina) for grants to perform molecular studies in the University of Colorado at Boulder, and IDEAWILD for the equipment provided for fieldwork.

## AUTHOR CONTRIBUTIONS

R.D. and S.D.S designed the study; R.D. and S.D.S extracted DNA and performed PCR; R.D. analyzed sequences, performed alignments and phylogenetic analyses; R.D. and M.D.L. applied phylogenetic comparative methods; R.D. and S.D.S. wrote the paper, with contributions from M.D.L. and G.E.B.

## DATA ACCESSIBILITY

All sequences have been deposited in GenBank (http://www.ncbi.nlm.nih.gov/genbank) with accessions numbers and voucher information detailed in Appendix S1. Gene trees are presented in Appendix S7.

## SUPPORTING INFORMATION

Additional Supporting Information may be found online in the supporting information tab for this article.

## Appendix 1.

Summary of taxon sampling, provenance, voucher (collector and number or barcode, in italics), herbarium where vouchers were housed between parenthesis (acronyms follow Index Herbariorum; Thiers, 2017), and GenBank accession numbers provided in the following order: ITS, LEAFY, *trnL*-*F*, *waxy*. ‘NA’ indicates either voucher or provenance information was not found, and ‘na’ that the region was not sampled for this accession. Newly generated sequences are indicated with an asterisk following the accession number.

***Alkekengi officinarum*** Mill. **var. *officinarum***, HUNGARY, cultivated, *ISZ 10*-*02*, na, na, HM006825, na. UNITED STATES, cultivated, *Whitson 1280* (DUKE), AY665850, na, na, na; NA, *D’Arcy 17707* (MO), na, MH822152^∗^, na, DQ169012. ***Alkekengi officinarum* var. *franchetii*** (Mast.) R.J.Wang, NA, *Lester S. XYZ* (BIRM), na, MH822151^∗^, MH752594^∗^, MH796557^∗^.

***Archiphysalis chamaesarachoides*** (Makino) Kuang, CHINA, Zhejiang, Gutian Mountain, *Li et al. 393* (HSNU), KC768877, na, KC768879, na.

***Aureliana angustifolia*** Alm.-Lafetá, BRAZIL, Minas Gerais, Juiz de Fora, *Giacomin et al. 965* (BHCB), KC832782, na, KC549633-KC549614, na. ***Aureliana anonacea*** (Sendtn.) I.M.C. Rodrigues & Stehmann (= *A. pereirae*), BRAZIL, Minas Gerais, Caraça Sanctuary, *Oliveira et al. 388* (BHCB), KC832788, na, KC549639-KC549620, na. BRAZIL, Mina Gerais, Caraça, *Barboza 3638b* (CORD), na, MH822153^∗^, na, KX690166. ***Aureliana brasiliana*** (Hunz.) Barboza & Hunz., BRAZIL, Rio de Janeiro, Itatiaia National Park, *Rodrigues et al. 106* (BHCB), KC832783, na, KC549634-KC549615, na. BRAZIL, Río de Janeiro, Petrópolis, *Barboza et al. 2055* (CORD), na, MH822154^∗^, na, MH796558^∗^. ***Aureliana cuspidata*** (Witasek) I.M.C. Rodrigues & Stehmann, BRAZIL, Sao Paulo, Conservation Area Boracéia, *Stehmann et al. 4812* (BHCB), KC832784, na, KC549635-KC549616, na. ***Aureliana darcyi*** Carvalho & Bovini, BRAZIL, Rio de Janeiro, Trindade, Paraty, *Stehmann et al. 4856* (BHCB), KC832785, na, KC549636-KC549617, na. ***Aureliana fasciculata*** (Vell.) Sendtn (= *A. fasciculata* var. *fasciculata*), BRAZIL, São Paulo, Jundiaí, Serra do Japi, *Stehmann et al. 4790* (BHCB), KC832786, na, KC549637-KC549618, na. BRAZIL, Paraná, Morretes, La Graciosa, *Barboza et al. 1630* (CORD), na, na, na, EF537144. ***Aureliana martiana*** (Sendtn.) I. M. C. Rodrigues & Stehmann, BRAZIL, Minas Gerais, Juiz de Fora, *Giacomin et al.* (BHCB), KC832787, na, KC549638-KC549619, na. ***Aureliana micrantha*** Sendtn., BRAZIL, Bahia, Road São José, *Stehmann 5064* (BHCB), KC832780, na, KC549631-KC549612, na. ***Aureliana picta*** (Mart.) I.M.C. Rodrigues & Stehmann, BRAZIL, São Paulo, Bananal, *Giacomin 887* (BHCB), KC832789, na, KC549640-KC549621, na. ***Aureliana pogogena*** (Moric.) I.M.C. Rodrigues & Stehmann, BRAZIL, Bahia, Conservation Area Serra Bonita, Camacan, *Stehmann 5084* (BHCB), KC832790, na, KC549641-KC549622, na. BRAZIL, *Stehmann et al. 5098* (BHBC), na, MH822155^∗^, na, MH796559^∗^. ***Aureliana sellowiana*** (Sendtn.) Barboza & Stehmann, BRAZIL, São Paulo, Parelheiros, *Rodrigues 69* (BHCB), KC832781, na, KC549632-KC549613, na. BRAZIL, São Paulo, desde Parelheiros rumbo a Eng. Marsilac, *Barboza et al. 2024* (CORD), na, MH822156^∗^, na, MH796560^∗^. ***Aureliana* sp. nov.** (= *A. fasciculata* var. *longifolia*), BRAZIL, São Paulo, Moji das Cruzes, *Stehmann 4800* (BHCB), KC832798, na, KC549649-KC549630, na. ***Aureliana tomentosa*** Sendtn. (= *A. fasciculata* var. *tomentella*), BRAZIL, Espiritu Santo, Santa Teresa, *Stehmann et al. 4857* (BHCB), KC832791, na, KC549642-KC549623, na. ***Aureliana velutina*** Sendtn. BRAZIL, Minas Gerais, Nova Lima, *Stehmann et al. 4543* (BHCB), KC832792, na, KC549643-KC549624, na. ***Aureliana wettsteiniana*** (Witasek) Hunz. & Barboza, BRAZIL, Santa Catarina, Porto União, *Thode 300* (BHCB), KC832793, na, KC549644-KC549625, na. BRAZIL, Paraná, Morretes, *Barboza 2020* (CORD), na, MH822157^∗^, na, MH796561^∗^.

***Brachistus nelsonii*** (Fernald) D’Arcy, J.L. Gentry & Averett, MEXICO, Campeche, Calakmul, Rancho El Sacrificio, *Martínez et al. 28097* (MEXU), MH763701^∗^, MH822158^∗^, MH752595^∗^, MH796562^∗^. **Brachistus stramonifolius** (Kunth) Miers, GUATEMALA, Solola and Chimaltenango, *Williams 41524* (DUKE), AY665845, na, na, na. MEXICO, Veracruz, Xalapa, Sierra Madre Oriental, *Sousa-Peña 738a* (MEXU), na, MH822159^∗^, EU580963, na.

***Calliphysalis carpenteri*** (Riddell) Whitson, UNITED STATES, Florida, *Whitson 1133* (DUKE), AY665851, MH822160^∗^, EU581042, MH796563^∗^.

***Capsicum lycianthoides*** Bitter, ECUADOR, Pichincha, Bellavista Cloud Forest Reserve, *Smith 203* (WIS), DQ314158, DQ309518, MH281754^∗^, DQ309468.

***Chamaesaracha arida*** Henrickson, UNITED STATES, New Mexico, Grant, San Vicente Creek drainage, *Deanna et al. 221* (COLO), MH763702^∗^, MH822161^∗^, MH752596^∗^, MH796564^∗^. ***Chamaesaracha coniodes*** (Moric. ex Dunal) Benth. & Hook. f. ex B.D. Jacks. & al., UNITED STATES, New Mexico, Harding, Ute Creek Valley, *Deanna et al. 234* (COLO), MH763703^∗^, MH822162^∗^, MH752597^∗^, MH796565^∗^. ***Chamaesaracha coronopus*** (Dunal) A. Gray, UNITED STATES, Colorado, Pueblo, Lake Pueblo, *Deanna & Carrasco 237* (COLO, CORD), MH763704^∗^, MH822163^∗^, MH752598^∗^, MH796566^∗^. ***Chamaesaracha crenata*** Rydb., MEXICO, Coahuila, Cepeda, Estación Marte, Talud norte, *Villarreal et al. 6646* (MEXU), MH763705^∗^, MH822164^∗^, MH752599^∗^, MH796567^∗^. ***Chamaesaracha pallida*** Averett, MEXICO, Zacatecas, Concepción del Oro, Sierra Astillero, *Villarreal & Ramírez 9391* (MEXU), MH763706^∗^, MH822165^∗^, MH752600^∗^, MH796568^∗^. ***Chamaesaracha rzedowskiana*** Hunz, MEXICO, Queretaro, Jalpan, Los Sarros, *López Ch. 546* (MEXU), MH763707^∗^, MH822166^∗^, MH752601^∗^, MH796569^∗^. ***Chamaesaracha sordida*** (Dunal) A. Gray, MEXICO, Sonora, Naco, Chihuahuan desert, *van Devender et al. 2003*-*352* (MEXU), MH763708^∗^, MH822167^∗^, MH752602^∗^, na. ***Chamaesaracha villosa*** Rydb., UNITED STATES, Texas, Pecos, Picnic Area East of Iraan, *Deanna et al. 211* (COLO), MH763709^∗^, MH822168^∗^, MH752603^∗^, MH796570^∗^.

***Cuatresia colombiana*** Hunz., COLOMBIA, Cauca, El Tampo, PNN Munchique, *Orozco et al. 3816* (COL, CORD), MH763710^∗^, MH822169^∗^, MH752604^∗^, MH796571^∗^. ***Cuatresia cuneata*** (Standl.) Bohs, NA, *Bohs 2394* (UT), MH763711^∗^, MH822170^∗^, MH752605^∗^, MH796572^∗^. ***Cuatresia cuspidata*** (Dunal) Hunz., COLOMBIA, Cundinamarca, Soacha, *Deanna 161* (CORD), MH763712^∗^, MH822171^∗^, MH752606^∗^, MH796573^∗^. ***Cuatresia exiguiflora*** (D’Arcy) Hunz., NA., Bohs 2454 (UT), MH763713^∗^, MH822172^∗^, EU580981, MH796574^∗^. ***Cuatresia foreroi*** Hunz., ECUADOR, Sucumbios, from Lumbaqui to La Bonita, *Croat & Ferry 93692* (MO), MH763714^∗^, na, na, na. ***Cuatresia fosteriana*** Hunz., NA, *Bohs 2753* (UT), MH763715^∗^, MH822173^∗^, MH752607^∗^, MH796575^∗^. ***Cuatresia garciae*** Hunz., COLOMBIA, Antioquia, Frontino, road to Murri, *Brant & Martínez 1410* (MO), na, MH822174^∗^, na, na. **Cuatresia plowmanii** Hunz., COLOMBIA, Bocayá, Santa María, Calichana, La Almenara, *Orejuela et al. 120* (COL), MH763716^∗^, MH822175^∗^, MH752608^∗^, MH796576^∗^. ***Cuatresia riparia*** (Kunth) Hunz., NA, *Bohs 2551* (UT), na, MH822176^∗^, EU580982, MH796577^∗^. ***Cuatresia* sp.**, ECUADOR, Pichincha, Bellavista Cloud Forest Reserve, *Smith 204* (WIS), DQ314165, DQ301518, KM200029, DQ309475. ***Cuatresia trianae*** Hunz. COLOMBIA, Caquetá, Florencia, corregimiento el Caraño, *Trujillo & Sánchez 3587* (HUAZ), MH763717^∗^, MH822177^∗^, MH752609^∗^, MH796578^∗^.

***Darcyanthus spruceanus*** (Hunz.) Hunz., PERU, Madre de Dios, Tambopata, Puerto Maldonado, *Valenzuela & Huamantupa 1011* (MO), na, na, MH752610^∗^, na.

***Deprea abra*-*patriciae*** (S. Leiva & Barboza) S. Leiva & Deanna, PERU, Amazonas, Bongará, Área de Conservación Privada Abra-Patricia, *Deanna & Leiva González 41* (CORD, HAO), KX557300, na, MH281755^∗^, KX690167. ***Deprea altomayoensis*** (S. Leiva & Quip.) Barboza & Deanna, PERU, San Martín, Rioja, Bosque de Protección Alto Mayo, *Deanna & Leiva González 84* (CORD), KX557302, MH822178^∗^, MH281756^∗^, KX690168. ***Deprea andersonii*** (N.W. Sawyer) Deanna & S. Leiva, ECUADOR, Napo, carretera Hollín-Loreto, km 26.5 (Ruta E45A, Troncal amazónica), *Deanna & Leiva González 116* (CORD, HAO), KX557301, MH822179^∗^, MH281757^∗^, KX690169. ***Deprea auccana*** S. Leiva, Barboza & Deanna, PERÚ, Amazonas, Bongará, Nueva Cajamarca – Pomacochas, *Deanna & Leiva González 44* (CORD), KX557303, MH822180^∗^, MH281758^∗^, KX690170. ***Deprea bitteriana*** (Werderm.) N.W. Sawyer & Benítez, COLOMBIA, Cundinamarca, Subachoque, El Tablazo, *Orozco et al. 3871* (COL, CORD), KP267794, MH822181^∗^, MH281760^∗^, KP267808. ***Deprea bongaraensis*** (S. Leiva) Deanna & Barboza, PERU, Amazonas, Bongará, carretera Bongará-Nuevo Cajamarca, *Deanna & Leiva González 36* (CORD), KX557304, MH822182^∗^, MH281761^∗^, KX690171. ***Deprea chotanae*** (S. Leiva, Pereyra & Barboza) S. Leiva, PERU, Cajamarca, Chota, bosque El Pargo, La Loma, *Deanna & Leiva González 59* (CORD), KX557305, MH822183^∗^, MH281762^∗^, KX690172. ***Deprea cuyacensis*** (N.W. Sawyer & S. Leiva) S. Leiva & Lezama, PERU, Piura, Ayabaca, bosque de Cuyas, *Barboza et al. 3367* (CORD), KP267793, MH822184^∗^, MH281763^∗^, KP267807. ***Deprea cyanocarpa*** Garzón & C.I. Orozco, COLOMBIA, *Muñoz 2* (COL), KP267797, MH822185^∗^, MH281764^∗^, KP267811. ***Deprea darcyana*** (N.W. Sawyer) Barboza & S. Leiva, COLOMBIA, Cauca, El Tambo, Parque Nacional Munchique, *Orozco et al. 3860* (COL, CORD), KX557306, na, MH281765^∗^, KX690173. ***Deprea ecuatoriana*** Hunz. & Barboza, ECUADOR, Zamora Chinchipe, Yanganá, rumbo al Cerro Toledo, *Orozco et al. 3952* (CORD), KP267795, MH822186^∗^, MH281767^∗^, KP267809. ***Deprea glabra*** (Standl.) Hunz., COLOMBIA, Cauca, El Tambo, Parque Nacional Munchique, *Orozco et al. 3812* (COL, CORD, QCA), KP267799, MH822187^∗^, MH281768^∗^, KP267813. ***Deprea harlingiana*** (Hunz. & Barboza) Deanna & S. Leiva, ECUADOR, Zamora Chinchipe, Parque Nacional Podocarpus, *Deanna & Leiva González 12* (CORD, HAO), KX557307, MH822188^∗^, MH281769^∗^, KX690174. ***Deprea hawkesii*** (Hunz.) Deanna, COLOMBIA, Cauca, El Tambo, Parque Nacional Munchique, *Orozco et al. 3824* (COL, CORD), KP267821, na, MH281770^∗^, KP267820. COLOMBIA, Huila, La Plata, Agua Bonita, Finca Meremberg, *Orejuela & Deanna 2568* (CORD, JBB), na, MH822189^∗^, na, na. ***Deprea longipedunculata*** (S. Leiva, E. Rodr. & J. Campos) Barboza, PERU, Cajamarca, San Ignacio, Tabaconas, caserío La Bermeja, *Deanna & Leiva González 18* (CORD, HAO), KX557309, MH822190^∗^, MH281775^∗^, KX690177. ***Deprea lutea*** (S. Leiva) Deanna, PERU, Cajamarca, Chota, km 46 desde desvío Llama-Huambos hacia La Granja, *Deanna & Leiva González 68* (CORD, HAO), KX557310, MH822191^∗^, MH281779^∗^, KX690178. ***Deprea macasiana*** (Deanna, S. Leiva & Barboza) Barboza, ECUADOR, Pastaza, Macas, cerro San José del Quílamo, *Deanna & Leiva González 111* (CORD, HAO, QUSF), KX557311, MH822192^∗^, MH281780^∗^, KX690180. ***Deprea maculatifolia*** (E. Rodr. & S. Leiva) S. Leiva, PERU, Amazonas, Bagua, Imaza, Comunidad Aguaruna de Yamayakat, *Deanna & Leiva González 82* (CORD, HAO), KX557313, na, MH281781^∗^, KX690181. ***Deprea micrantha*** S. Leiva & Barboza, ECUADOR, Zamora Chinchipe, Reserva Biológica San Francisco, Leiva González & Barboza 6530 (CORD, HAO, LOJA), MH281823^∗^, na, MH281776^∗^, MH281832^∗^. ***Deprea nieva*** (S. Leiva & N.W. Sawyer) Barboza & Deanna, PERU, Amazonas, Bongará, km 384, bordes de carretera Nueva Cajamarca-Pomacochas (Florida), *Deanna & Leiva González 46* (CORD, HAO), KP267769, MH304887^∗^, MH281782^∗^, KP267763. ***Deprea nubicola*** N.W. Sawyer, COLOMBIA, Magdalena, Ciénaga, Sierra Nevada de Santa Marta, *Orejuela & Vélez 215* (COL), KP267796, MH822193^∗^, MH281783^∗^, KP267810. ***Deprea orinocensis*** (Kunth) Raf., VENEZUELA, *Benítez & Mancilla 7460* (MY), KP267767, MH822194^∗^, MH281784^∗^, KP267762. ***Deprea paneroi*** Benítez & M. Martínez, VENEZUELA, *Benítez et al. 7454* (MY), KP267768, na, MH281785^∗^, KP267761. ***Deprea parviflora*** (N.W. Sawyer & S. Leiva) S. Leiva, PERÚ, Cajamarca, Cutervo, km 1543-1544, carretera Cutervo-La Capilla, *Deanna & Leiva González 73* (CORD, HAO), KX557314, MH822195^∗^, MH281786^∗^, KX690183. ***Deprea pauciflora*** Deanna, Barboza & S. Leiva, ECUADOR, Zamora Chinchipe, límite del Parque Nacional Podocarpus, *Deanna & Leiva González 13* (CORD), KX557332, MH822196^∗^, MH281787^∗^, KX690182. ***Deprea pecaensis*** S. Leiva, Deanna & Barboza, PERU, Amazonas, Bagua, La Peca, puente El Arenal, *Deanna & Leiva González 49* (CORD, HAO), KX557315, MH822197^∗^, MH281789^∗^, KX690184. ***Deprea pedrazae*** (S. Leiva & Barboza) Deanna & S. Leiva, PERU, Amazonas, Bagua, La Peca, puente El Arenal, *Deanna & Leiva González 48* (CORD, HAO), KX557316, MH822198^∗^, MH281788^∗^, KX690185. ***Deprea physalidicalyx*** S. Leiva, Barboza & Deanna, PERU, San Martín, San Martín, carretera Tarapoto hacia Bella Vista, *Leiva González & Barboza 5645* (CORD, HAO), KX557341, MH822199^∗^, MH281790^∗^, KX690186. ***Deprea pilosa*** (S. Leiva, E. Rodr. & J. Campos) Deanna, PERU, Cajamarca, San Ignacio, San José de Lourdes, Estrella del Oriente, *Deanna & Leiva González 32* (CORD, HAO), KX557317, MH822200^∗^, MH281791^∗^, KX690187. ***Deprea pomacochaensis*** (S. Leiva) Barboza, PERU, Amazonas, Bongará, carretera Bongará-Nueva Cajamarca, *Deanna & Leiva González 33* (CORD, HAO), KX557318, MH822201^∗^, MH281792^∗^, KX690188. ***Deprea psilophyta*** (N.W. Sawyer) S. Leiva & Deanna, ECUADOR, Loja, Nudo de Sabanilla, sendero a Ayupallas, *Orozco et al. 3947* (COL, CORD), na, na, MH281793^∗^, na. ECUADOR, Zamora Chinchipe, carretera desde Yanganá hacia Valladolid, *Sawyer 770* (CONN, LOJA), KP267772, na, na, KP267766. ***Deprea pumila*** (S. Leiva, Barboza & Deanna) S. Leiva, ECUADOR, Pastaza, Mera, camino al río Anzú, *Orozco et al. 3890* (COL, CORD, QCA), KX557320, MH304886^∗^, MH281794^∗^, KX690189. ***Deprea purpurea*** (S. Leiva) Barboza & S. Leiva, PERU, Cajamarca, San Ignacio, San José de Lourdes, Estrella del Oriente, *Deanna & Leiva González 27* (CORD, HAO), KX557319, MH822202^∗^, MH281795^∗^, KX690192. ***Deprea purpureocarpa*** (S. Leiva, Deanna & Barboza) Deanna, ECUADOR, Napo, carretera Cosanga-Baeza, 5.4 km al sur de Baeza, Deanna & Leiva González 125 (CORD, HAO, QCNE), KX557321, MH822203^∗^, MH281800^∗^, KX690193. ***Deprea sachapapa*** (Hunz.) S.
Leiva & Deanna, ECUADOR, Cotopaxi, San Francisco de las Pampas, Otonga, *Orozco et al. 3985* (COL, CORD, QCA), KX557328, na, MH281796^∗^, KX690197. ECUADOR, Pichincha, *Smith 205* (WIS), na, DQ301519, na, na. ***Deprea sagasteguii*** (S. Leiva, Quip. & N.W. Sawyer) Barboza, PERU, Piura, Ayabaca, cerro Aypate, *Deanna & Leiva González 97* (CORD, HAO), KX557330, MH822204^∗^, MH281797^∗^, KX690200. ***Deprea sapalachensis*** S. Leiva & Barboza, PERU, Piura, Huancabamba, Carmen de la Frontera, *Barboza & Leiva González 4833* (CORD, HAO), na, na, MH752611^∗^, MH796579^∗^. ***Deprea sawyeriana*** (S. Leiva, E. Rodr. & J. Campos) S. Leiva, PERU, Cajamarca, San Ignacio, Tabaconas, caserío La Bermeja, *Deanna & Leiva González 14* (CORD, HAO), KX557331, MH822205^∗^, MH281798^∗^, KX690202. ***Deprea* sp.**, ECUADOR, Pastaza, Mera, desde la Plaza Mayor de Mera hacia Cavernas del Río Anzú, *Deanna et al. 114* (CORD), MH763718^∗^, na, MH752612^∗^, na. ***Deprea steyermarkii*** (Hunz.) S. Leiva & Barboza, ECUADOR, Azuay, carretera Gualaceo-Indanza, km 23, *Deanna & Leiva González 108* (CORD, HAO), KX557335, MH822206^∗^, MH281803^∗^, KX690203. ***Deprea subtriflora*** (Ruiz & Pav.) D’Arcy, BOLIVIA, La Paz, Nor-Yungas, carretera desde Chuspipata a Coroico, *Barboza & Leiva González 3663* (CORD), KP267770, MH822207^∗^, MH281805^∗^, KP267764. ***Deprea sylvarum*** (Standl. & C.V. Morton) Hunz., COSTA RICA, *Bohs 2504* (UT), KP267800, na, MH281806^∗^, KP267814. ***Deprea teresitae*** Deanna & Orejuela, COLOMBIA, Valle del Cauca, Reserva ‘El Refugio’, *Deanna & Calderón 169* (PSO, CORD), MH281825^∗^, na, MH281801^∗^, MH281833^∗^. ***Deprea toledoana*** (Barboza & S. Leiva) Barboza, ECUADOR, Zamora Chinchipe, a Valladolid desde Yanganá, *Orozco et al. 3936* (COL, CORD, QCA), KX557337, MH822208^∗^, MH281807^∗^, KX690205. ***Deprea vasquezii*** (S. Leiva, E. Rodr. & J. Campos) Deanna, PERU, Cajamarca, San Ignacio, San José de Lourdes, Estrella del Oriente, *Deanna & Leiva González 28* (CORD, HAO), KX557339, MH822209^∗^, MH281808^∗^, KX690207. ***Deprea zakii*** Barboza, S. Leiva & Deanna, ECUADOR, Napo, Quijos, carretera Papallacta-Cuyuja, *Deanna et al. 138* (CORD, QCNE), KX557340, MH822210^∗^, MH281802^∗^, KX690208. ***Deprea zamorae*** Barboza & S. Leiva, ECUADOR, Zamora Chinchipe, Parque Nacional Podocarpus, *Orozco et al. 3926* (COL, CORD, QCA), KP267792, MH822211^∗^, MH281809^∗^, KP267806.

***Discopodium penninervium*** Horchst., TANZANIA, *Tanner 3288*, KC832794, MH822212^∗^, na, na. UGANDA, Kabarole, Burahya, *Knapp 9808* (BM), na, na, EU580986, na.

***Dunalia brachyacantha*** Miers, ARGENTINA, Jujuy, Valle Grande, *Nee & Bohs 50811* (NY), DQ314172, DQ301527, MH281810^∗^, DQ309482. ***Dunalia obovata*** (Ruiz & Pav.) Dammer, PERU. Junin, *Smith et al. 458* (HAO, F, MO, NY, USM, WIS), DQ314192, DQ301547, MH281811^∗^, MDQ309499. ***Dunalia spathulata*** Ruiz & Pav.) Braun & Bouché, PERU, Huanuco, *Smith et al. 452* (HAO, F, MO, NY, USM, WIS), DQ314198, DQ301554, MH752613^∗^, DQ309506. ***Dunalia spinosa*** (Meyen) Dammer, BOLIVIA, Potosí, Tomas Frias, *Smith et al. 379* (MO, WIS) DQ314188, DQ301543, MH281812^∗^, DQ309495.

***Eriolarynx fasciculata*** (Miers) Hunz., BOLIVIA, Cochabamba, *Smith et al. 432* (HAO, F, MO, NY, WIS), DQ314196, DQ301552, MH752614^∗^, DQ309504. ***Eriolarynx iochromoides*** (Hunz.) Hunz., ARGENTINA, Catamarca, Andalgalá, Río Potrero, *Barboza et al. 1966* (CORD), KP267802, MH304888^∗^, MH281813^∗^, KP267816. ***Eriolarynx australis*** (Griseb.) J.M.H Shaw, BOLIVIA, Chuquisaca, *Smith et al. 390* (WIS), DQ314189, DQ301544, KP756712, DQ309496. ***Eriolarynx lorentzii*** (Dammer) Hunz., ARGENTINA, Tucumán, *Hawkes et al. 3452* (BIRM), DQ314171, DQ301525, KP756713, DQ309481.

***Iochroma amicorum*** M. Cueva, S.D. Sm. & S. Leiva, PERU, Oxapampa, Huancabamba, PN Yanachaga-Chemillen, *Smith 542* (HAO, HOXA, MO, USM), KM514683, KM514684, MH752615^∗^, KM521199. ***Iochroma arborescens*** (L.) J.M.H. Shaw, COSTA RICA, Puntarenas, Las Cruces, *Bohs 2428* (UT), DQ314173, DQ301528, KP756700, DQ309483. ***Iochroma ayabacense*** S. Leiva, PERU, Piura, Ayabaca, *Smith & Leiva González 337A* (HAO, F, MO, WIS), DQ314194, DQ301549, MH752616^∗^, DQ309501. ***Iochroma barbozae*** S. Leiva & Deanna, PERU, Piura, Ayabaca, *Deanna et al. 91* (CORD), MH763719^∗^, MH822213^∗^, MH752617^∗^, MH796581^∗^. ***Iochroma baumii*** S.D. Sm. & S. Leiva, ECUADOR, Napo, Papallacta, *Smith & López 476* (QCNE, F, WIS), DQ314202, DQ301558, MH752618^∗^, DQ309513. ***Iochroma calycinum*** Benth., ECUADOR, Pichincha, *Smith 471* (F, QCNE, WIS), DQ314201, DQ301557, MH281815^∗^, DQ309512. ***Iochroma confertiflorum*** (Miers) Hunz., ECUADOR, Loja, *Smith et al. 237* (QCNE, MO, WIS), DQ314176, DQ301531, MH752619^∗^, DQ309486. ***Iochroma cornifolium*** (Kunth) Miers, ECUADOR, Loja, *Smith et al. 242* (QCNE, MO, WIS), DQ314177, DQ301532, MH752620^∗^, DQ309487. ***Iochroma cyaneum*** (Lindl.) G.H.M. Lawr. & J.M. Tucker, ECUADOR, Loja, Catamayo-El Cisne road, *Smith 223* (QCNE, MO, WIS), DQ314180, DQ301535, MH281814^∗^, DQ309490. ***Iochroma edule*** S.Leiva, PERU, La Libertad, *Smith et al. 300* (HAO, F, MO, NY, USM, WIS), DQ314193, DQ301548, KP756703, DQ309500. ***Iochroma ellipticum*** (Hook. f.) Hunz., ECUADOR, Galápagos, *Jager 622* (CDS), DQ314199, DQ301555, MH752622^∗^, DQ309507. ***Iochroma fuchsioides*** (Bonpl.) Miers, ECUADOR, Azuay, *Smith & López 488* (QCNE, F, MO, WIS), DQ314203, DQ301559, KP756711, DQ309514. ***Iochroma gesnerioides*** (Kunth) Miers, ECUADOR, Pichincha, Reserva Geobotanica Pululahua, *Smith 200* (QCNE, MO, WIS), DQ314179, DQ301534, MH281816^∗^, DQ309489. ***Iochroma lehmannii*** Dammer ex Bitter, ECUADOR, Cañar, *Smith & López 484* (QCNE, F, MO, WIS), DQ314200, DQ301556, MH752623^∗^, DQ309511. ***Iochroma loxense*** (Kunth) Miers, ECUADOR, Loja, *Smith 220* (QCNE, MO, WIS), DQ314175, DQ301530, MH752624^∗^, DQ309485. ***Iochroma nitidum*** S. Leiva & Quip., PERU, Amazonas, *Smith et al. 371* (HAO, F, MO, NY, USM, WIS), DQ314168, DQ301521, MH752625^∗^, DQ309478. ***Iochroma parvifolium*** (Roem. & Schult.) D’Arcy, PERU, La Libertad, *Smith et al. 303* (HAO, F, MO, NY, USM, WIS), DQ314195, DQ301551, MH752626^∗^, DQ309503. ***Iochroma peruvianum*** (Dunal) J.F. Macbr., PERU, Cajamarca, *Smith & Hall 353* (HAO, F, MO, NY, USM, WIS), DQ314197, DQ301553, KP756706, DQ309505. ***Iochroma piuranum*** S.Leiva, PERU, Piura, Ayabaca, *Deanna et al. 93* (CORD), MH763721^∗^, MH822215^∗^, MH752627^∗^, MH796582^∗^. ***Iochroma salpoanum*** S. Leiva & Lezama, PERU, La Libertad, ***Smith 364*** (WIS), DQ314187, DQ301542, MH752628^∗^, DQ309509. ***Iochroma squamosum*** S. Leiva & Quip., PERU, Piura, Ayabaca, *Smith et al. 330* (HAO, F, MO, NY), DQ314186, DQ301541, MH281817^∗^, DQ309494. ***Iochroma stenanthum*** S. Leiva, Quip. & N.W. Sawyer, PERU, Cajamarca, *Smith et al. 313* (HAO, F, MO, NY, USM, WIS), DQ314184, DQ301539, MH752629^∗^, DQ309508. ***Iochroma tingoanum*** S. Leiva, PERU, Amazonas, *Smith et al. 370* (HAO, F, MO, NY, USM, WIS), DQ314167, DQ301520, MH752630^∗^, DQ309477. ***Iochroma tupayachianum*** S. Leiva, PERU, La Libertad, *Smith et al. 526* (F, MO, USM, WIS), KC290442, KC290441, na, KC243428.

***Leucophysalis grandiflora*** (Hook.) Rydb., UNITED STATES, *Olmstead S*-*30* (WTU), DQ314162, DQ301515, EU581013, DQ309472. ***Leucophysalis nana*** (A. Gray) Averett, UNITED STATES, *Bartholomew 5994* (MO), MH763722^∗^, na, EU581014, na.

***Lycianthes inaequilatera*** Bitter, ECUADOR, Pichincha, Alluriquin, *Smith 210* (WIS), DQ314159, DQ309519, na, DQ309469. BOLIVIA, *Bohs 3089* (UT), na, na, EU581018, na.

***Nothocestrum breviflorum*** A. Gray, HAWAII, Hamakua, Kailikaula Cliffs and Stream, *Wood et al. 4862* (MO), MH763723^∗^, MH822216^∗^, MH752631^∗^, MH796583^∗^.

***Nothocestrum latifolium*** A. Gray, HAWAII, Polynesia Hawaiian Islands, *H. St. John 24469*, KC832796, na, na, na. HAWAII, *Herbst et al. 725* (COLO), na, na, EU581037, na. HAWAII,
*Lorentz 9063*, na, MH822217^∗^, na, MH796584^∗^. ***Nothocestrum longifolium*** A. Gray, HAWAII, Main Hawaiian Islands, North Hilo, *Cuddihy 743* (BISH), KC832795, na, na, na. HAWAII, *Oppenheimer s.n.* (BISH), na, MH822218^∗^, EU581038, MH796585^∗^. ***Nothocestrum peltatum*** Skottsb., HAWAII, Honopu, NW of Kainamanu, Acacia koa, *Wood & Query 15166* (MO), na, MH822219^∗^, MH752632^∗^, MH796586^∗^.

***Oryctes nevadensis*** S. Watson, UNITED STATES, Nevada, Churchill, *Tiehm 11982* (COLO, TEX), AY665864, na, EU581039, na.

***Physaliastrum echinatum*** (Yatabe) Makino, CHINA, Yunyougu, Xinchengzi Town, Miyun District, Beijing, *Liu & Shi 5186* (PE), MH763724^∗^, MH822220^∗^, MH752633^∗^, MH796587^∗^. ***Physaliastrum japonicum*** (Franch. & Sav.) Honda, NA, *YYZWF20387*, KP894015, na, na, na. ***Physaliastrum heterophyllum*** (Hemsl.) Migo, CHINA, Zhejiang West Tianmu Mountain, *Li et al. 435* (HSNU), KC768878, na, KC768880, na. ***Physaliastrum sinense*** (Hemsl.) D’Arcy & Z.Y. Zhang, CHINA, Sichuan, *Hunggui 1177* (MO), na, MH822221^∗^, na, na.

***Physalis acutifolia*** (Miers) Sandwith, UNITED STATES, Arizona, Cpcjose, *Makings 3742* (MO), na, MH822222^∗^, MH752634^∗^, MH796588^∗^. UNITED STATES, cultivated, *NIJ 974750059*, AY665876, na, na, na. ***Physalis angulata*** L., ARGENTINA, Córdoba, Río Seco, Ruta Nac. Nº 9, pasando Va. de María, *Morero 365* (CORD), MH763725^∗^, MH822223^∗^, MH752635^∗^, MH796589^∗^. ***Physalis angustifolia*** Nutt., UNITED STATES, Florida, Okalossa, *Miller et al. 9107* (MO), na, MH822224^∗^, MH752636^∗^, na. UNITED STATES, Florida, *Whitson, no voucher*, AY665878, na, na, na. ***Physalis angustiphysa*** Nutt., MEXICO, Chiapas, *Ton 9286* (TEX), AY665879, na, na, na. ***Physalis arenicola*** Kearney, UNITED STATES, Florida, Putnam, Ordway-Swisher Biological Station, *Majure et al. 5075* (FLAS), na, MH822225^∗^, MH752637^∗^, MH796590^∗^. UNITED STATES, Florida, *Whitson, no voucher*, AY665880, na, na, na. ***Physalis campanula*** Standl. & Steyerm., MEXICO, Veracruz, *Ventura 4882* (MO), AY665882, na, na, na. ***Physalis campechiana*** L., MEXICO, Tamaulipas, *Jimenez 454* (TEX), AY665867, MH822226^∗^, MH752638^∗^, MH796591^∗^. ***Physalis caudella*** Standl., MEXICO, Chihuahua, *Quintana 3075* (TEX), AY665891, na, na, na. ***Physalis chenopodifolia*** Lam., MEXICO, México, Pirámides de Teotihuacan, *Chiarini et al. 1277* (CORD), na, MH304893^∗^, MH752639^∗^, MH304879^∗^. UNITED STATES, cultivated, *Whitson 1287* (DUKE), AY665883, na, na, na. ***Physalis cinerascens*** (Dunal) Hitchc. **var. *cinerascens***, UNITED STATES, Texas, Comal, Schmucks and Doeppens, roadsides, *Deanna et al. 206* (COLO, CORD), MH763726^∗^, MH822227^∗^, MH752640^∗^, MH796592^∗^. ***Physalis cinerascens* var. *spathulifolia*** (Torr.) J.R. Sullivan, UNITED STATES, Texas, Colorado, East to the Attwater Prairie Chicken National Wildlife Refuge, *Deanna et al. 203* (COLO), MH763727^∗^, MH822228^∗^, MH752641^∗^, MH796593^∗^. ***Physalis cordata*** Mill., PERU, Cajamarca, Contumazá, *Knapp et al. 10557* (CORD), MH763728^∗^, MH822229^∗^, MH752642^∗^, MH796594^∗^. ***Physalis coztomatl*** Dunal, MEXICO, *Garcia 264* (MO), AY665887, na, na, na. ***Physalis crassifolia*** Benth., UNITED STATES, California, *Sharples 744* (COLO), MH763729^∗^, MH822230^∗^, MH752643^∗^, MH796595^∗^. ***Physalis* x *elliottii*** Kunze, UNITED STATES, Florida, Sanibel Island, Bailey Tract, *Wheeler 14144* (SI), na, MH822231^∗^, MH752644^∗^, MH796596^∗^. ***Physalis fendleri*** A. Gray, UNITED STATES, New Mexico, Grant, outside of Silver City, *Deanna et al. 219* (COLO), MH763730^∗^, MH822232^∗^, MH752645^∗^, MH796597^∗^. ***Physalis glabra*** Benth., MEXICO, Baja California Sur, La Paz, *Provance et al. 8003* (MO), MH763731^∗^, na, MH752646^∗^, na. ***Physalis glutinosa*** Schlecht., MEXICO, Durango, *Sikes 375* (TEX), AY665892, na, na, na. ***Physalis greenmanii*** Waterf., MEXICO, Veracruz, *Nee 22432* (MO), AY665893, na, na, na. MEXICO, Veracruz, Villa Aldama, *Nee 32880* (CORD), na, MH822233^∗^, na, na. ***Physalis grisea*** (Waterf.) M. Martínez, UNITED STATES, cultivated, *NIJ 894750256*, AY665915, na, na, na. ***Physalis hederifolia*** A. Gray, UNITED STATES, Texas, Uvalde, dry Frio River, *Deanna et al. 209* (COLO), MH763732^∗^, MH822234^∗^, MH752647^∗^, MH796598^∗^. ***Physalis heterophylla*** Nees, UNITED STATES, Colorado, Larimer, Lory State Park, *Deanna et al. 199* (COLO), na, MH822235^∗^, na, MH796599^∗^. UNITED STATES, North Carolina, Caswell, *Whitson, no voucher*, AY665907, na, na, na. UNITED STATES, *Olmstead S-64* (WTU), na, na, EU581043, na. ***Physalis hintonii*** Waterf., MEXICO, Nuevo Leon, *Villarreal 4909* (MO), AY665895, na, na, na. ***Physalis ignota*** Britton, MEXICO, Chiapas, *Breedlove 52891* (MO), AY665897, na, na, na. ***Physalis ixocarpa*** Brot. ex Hornem., UNITED STATES, cultivated, *Deanna 251* (CORD), MH763733^∗^, MH822236^∗^, MH752648^∗^, MH796600^∗^. ***Physalis lagascae*** Roem. & Schult., PERU, Cajamarca, Cutervo, *Särkinen 4548* (BM), na, MH304892^∗^, MH752649^∗^, MH304880^∗^. ***Physalis lanceolata*** Michx., UNITED STATES, North Carolina, Scotland, *Horn 1133* (DUKE), AY665899, na, na, na. ***Physalis lassa*** Stand. & Steyerm., MEXICO, Comala, *Sanders 11807* (MO), AY665900, na, na, na. ***Physalis longifolia*** Nutt., UNITED STATES, New Mexico, Bernalillo, Albuquerque, *Deanna et al. 227* (COLO), na, MH822237^∗^, MH752650^∗^, MH796601^∗^. UNITED STATES, Kansas, Riley, *Whitson s.n.* (DUKE 358627), AY665901, na, na, na. ***Physalis macrosperma*** ined. UNITED STATES, Arkansas, Miller, *Gentry & Reid 3188*, MH763734^∗^, MH822238^∗^, MH752651^∗^, na. ***Physalis melanocystis*** (B.L. Rob.) Bitter, MEXICO, Tamaulipas, *Martinez 1940* (MO), AY665865, MH822239^∗^, MH752652^∗^, MH796602^∗^. ***Physalis microcarpa*** Urb., MEXICO, Chihuahua, *Laferriere 1661* (MO), AY665903, na, na, na. ***Physalis microphysa*** A. Gray, MEXICO, Coahuila, *Henrickson 11850* (TEX), AY665859, MH822240^∗^, MH752653^∗^, MH796603^∗^. ***Physalis minima*** L., AUSTRALIA, South Australia, cultivated, *Symon 14813* (CORD), na, MH822241^∗^, na, na. THAILAND, cultivated, *NIJ 974750167*, AY665904, na, na, na. ***Physalis minimaculata*** Waterf., MEXICO, Oaxaca, *Mayfield 986* (TEX), AY665906, na, na, na. ***Physalis mollis*** Nutt., UNITED STATES, Texas, Bastrop, *Deanna et al. 201* (COLO), na, MH822242^∗^, MH752654^∗^, MH796604^∗^. ***Physalis neomexicana*** Rydb., UNITED STATES, New Mexico, Santa Fe, *Deanna et al. 228* (COLO), MH763735^∗^, MH822243^∗^, MH752655^∗^, MH796605^∗^. ***Physalis nicandroides*** Schltdl., MEXICO, Veracruz, Acultzingo, *Nee 33132* (CORD), na, MH822244^∗^, na, MH796606^∗^. MEXICO, Morelos, *Hernandez 2488* (MO), AY665912, na, na, na. ***Physalis orizabae*** Dunal, MEXICO, Morelos, Lagunas de Zempoala, *Chiarini et al. 1280* (CORD), MH763736^∗^, MH822245^∗^, MH752656^∗^, MH796607^∗^. ***Physalis patula*** Mill., MEXICO, Ciudad de México, *Chiarini et al. 1273* (CORD), na, MH822246^∗^, MH752657^∗^, MH796608^∗^. MEXICO, Veracruz, *Nee 32810* (MO), AY665913, na, na, na. ***Physalis peruviana*** L., ECUADOR, Pichincha, cultivated, *Smith 217* (WIS), DQ314161, DQ301514, na, DQ309471. PERU, *Olmstead S-69* (WTU), na, na, EU581044, na. ***Physalis philadelphica*** Lam., UNITED STATES, cultivated, *Bohs 2433* (UT), na, MH822247^∗^, EU581045, MH796609^∗^. UNITED STATES, cultivated, *Whitson s.n.* (DUKE), AY665871, na, na, na. ***Physalis pruinosa* var. *argentina*** J. M. Toledo & Barboza, ARGENTINA, Jujuy, Valle Grande, Ruta P.N. Calilegua-San Francisco-Valle Grande, *Smith & Chiarini 630* (COLO), na, MH822248^∗^, MH752658^∗^, MH796610^∗^. ***Physalis pubescens* L. var. *pubescens***, MEXICO, Morelos, Lagunas de Zempoala, *Chiarini et al. 1281* (CORD), na, MH304895^∗^, MH752659^∗^, MH304881^∗^. COSTA RICA, La Selva Biological Station, *Whitson 3* (DUKE), AY665916, na, na, na. ***Physalis pubescens* var. *higrophyla*** (Mart.) Dunal, ARGENTINA, Jujuy, Ledesma, Libertador Gral. San Martín, *Toledo 1652* (CORD), MH763737^∗^, MH822249^∗^, MH752660^∗^, MH796611^∗^. ***Physalis pumila Nutt.* var. *pumila***, UNITED STATES, New Mexico, San Miguel, Sangre de Cristo Mountains, *Deanna et al. 230* (COLO), MH763738^∗^, MH822250^∗^, MH752661^∗^, MH796612^∗^. ***Physalis pumila* var. *hispida*** (Waterf.) W.F. Hinton, UNITED STATES, Colorado, Larimer, next to Poudre River, *Deanna et al. 200* (COLO), MH763739^∗^, MH822251^∗^, MH752662^∗^, MH796613^∗^. ***Physalis purpurea*** Wiggins, BOLIVIA, La Paz, Sud-Yungas, *Barboza 3657* (CORD), MH763740^∗^, MH822252^∗^, MH752663^∗^, MH796614^∗^. ***Physalis solanacea*** (Schltdl.) Axelius, MEXICO, Tamaulipas, Llera de Canales, *Nee & Calzada 33199* (CORD), na, MH822253^∗^, na, MH796615^∗^. MEXICO, cultivated, *Olmstead S*-*37* (WTU), AY665877, na, EU581025, na. ***Physalis sordida*** Fernald, MEXICO, Nuevo Leon, *Hinton 18464* (TEX), AY665869, na, na, na. ***Physalis subilsiana*** J.M. Toledo ARGENTINA, Salta, General José de San Martín, *Toledo & Domínguez 226* (CORD), na, MH822254^∗^, na, na. ***Physalis sulphurea*** (Fernald) Waterf., MEXICO, 1 km al N de San Juan Citlaltepec, *Rodríguez García 116* (CORD), MH763741^∗^, na, na, na. ***Physalis victoriana*** J.M. Toledo, ARGENTINA, Jujuy, Ruta Provincial Nº 1, de Caimancito a Palma Sola, *Carrizo García 5* (CORD), MH763742^∗^, na, MH752664^∗^, MH796616^∗^. ***Physalis virginiana*** Mill., UNITED STATES, Colorado, Boulder, *Deanna & Smith 238* (COLO), MH763743^∗^, MH822255^∗^, MH752665^∗^, MH796617^∗^. ***Physalis viscosa*** L., ARGENTINA, Córdoba, Calamuchita, *Deanna & Tamborini 179* (CORD), na, MH304894^∗^, MH752666^∗^, MH304882^∗^. UNITED STATES, cultivated, *Whitson 1282* (DUKE), AY665870, na, na, na. ***Physalis walteri*** Nutt., UNITED STATES, Florida, Levy, Havens Island, *Majure 3051* (FLAS), na, MH822256^∗^, MH752667^∗^, MH796618^∗^. UNITED STATES, Florida, *Whitson, no voucher*, AY665918, na, na, na.

***Quincula lobata*** (Torr.) Raf., UNITED STATES, New Mexico, Harding, *Deanna et al. 235* (COLO), MH763744^∗^, MH822257^∗^, MH752668^∗^, MH796619^∗^.

***Salpichroa tristis*** Miers, BOLIVIA, Potosí, Tomas Frias, *Smith et al. 382* (HAO, F, MO, NY, WIS), DQ314160, DQ309520, MH281774^∗^, DQ309470.

***Saracha andina*** Rob. Fernandez, I. Revilla & E. Pariente, PERU, Ayacucho, Lucanas, *Smith & Fernandez 594* (COLO, F, MO, USM), KY172041, KY172040, na, KY172039. ***Saracha nigribaccata*** J.M.H. Shaw, ECUADOR, Pichincha, *Smith 211A* (QCNE, MO, WIS), DQ314174, DQ301529, EU580988, DQ309484. ***Saracha punctata*** Ruiz & Pav., BOLIVIA,
La Paz, Nor Yungas, Rio Unduavi, *Nee 51804* (MO, NY), DQ314182, DQ301537, KP756709, DQ309492. ***Saracha quitensis*** (Hook.) Miers, ECUADOR, Napo, Laguna de Papallacta, *Smith 257* (QCNE, MO, WIS), DQ314178, DQ301533, MH281777^∗^, DQ309488.

***Schraderanthus viscosus*** (Schrad.) Averett, MEXICO, Oaxaca, *Torres 7932* (MO), AY665848, na, na, na.

***Trozelia grandiflora*** (Benth.) J.M.H. Shaw, PERU, Cajamarca, *Smith et al. 320A* (HAO, F, MO, NY, USM, WIS), DQ314170, DQ301523, MH752669^∗^, DQ309480. ***Trozellia umbellata*** (Ruiz & Pav.) Raf., PERÚ, La Libertad, *Smith et al. 301* (HAO, F, NY, USM, WIS), DQ314169, DQ301522, MH281818^∗^, DQ309479.

***Tubocapsicum anomalum***, CHINA, *Chen 231* (MO), DQ314163, DQ301516, EU581066, DQ309473.

***Tzeltalia amphitricha*** (Bitter) E. Estrada & M. Martínez, MEXICO, Chiapas, *Martínez 20523* (TEX), AY665853, na, na, na. ***Tzeltalia calidaria*** (Standl. & Steyerm.) E. Estrada & M. Martínez, GUATEMALA, *Lundell 19625* (TEX), na, na, MH752670^∗^, na. ***Tzeltalia esenbeckii*** M. Martínez & O. Vargas, MEXICO, Chiapas, La Independencia, from Las Margaritas to Campo Alegre, *Breedlove 51325* (MEXU), MH763745^∗^, na, MH752671^∗^, na.

***Vassobia breviflora*** (Sendtn.) Hunz., BOLIVIA, Chuquisaca, *Smith 412* (WIS), DQ314190, DQ301545, MH281819^∗^, DQ309497. ***Vassobia dichotoma*** (Rusby) Bitter, BOLIVIA, *Nee et al. 51797* (UT), na, na, EU581067, na. BOLIVIA, La Paz, *Smith 440* (WIS), DQ314191, DQ301546, na, DQ309498.

***Withania adpressa*** Cors., MORROCCO, *Lewalle 13205* (MO), na, na, MH752672^∗^, MH796620^∗^. ***Withania aristata*** Pauq., SPAIN, Canary Islands, *del Arco s.n.* (CORD), MH763746^∗^, MH822258^∗^, MH752673^∗^, MH796621^∗^. ***Withania coagulans*** (Stocks) Dunal, CENTRAL ASIA, *Olmstead S*-*109* (WTU), na, MH822259^∗^, EU581068, MH796622^∗^. ***Withania frutescens*** (L.) Pauquy, MOROCCO, Beldevere de Chicht, 15 km N of Essaouira, *Miller et al. 335* (MO), na, MH822260^∗^, na, na. ***Withania riebeckii*** Balf. f., NA., *D’Arcy 17750* (MO), na, MH822261^∗^, KC549645-KC549626, MH796623^∗^. ***Withania somnifera*** (L.) Dunal, NA, *Whitson 1262* (KNK), na, MH304890^∗^, na, MH304884^∗^. NA, *Lester S. 0960*, KC832797, na, na, na. SPAIN, Canary Is., Mediterranean to Central Asia, *Whitson s.n.* (KNK), na, na, EU581069, na.

***Witheringia asterotricha*** (Standl.) Hunz., COSTA RICA, *Bohs 3007* (UT), MH763747^∗^, MH822262^∗^, MH752674^∗^, MH796624^∗^. ***Witheringia coccoloboides*** (Dammer) Hunz. COSTA RICA, Bohs 2568 (UT), MH281826^∗^, MH304889^∗^, MH752675^∗^, MH304885^∗^. ***Witheringia correana*** D’Arcy, PANAMA, Bocas del Toro, Fortuna, *D’Arcy 16415* (MO), MH763748^∗^, MH822263^∗^, MH752676^∗^, MH796625^∗^. ***Witheringia killipiana*** Hunz., COLOMBIA, Cauca, El Tambo, *Orozco et al. 3858* (COL, CORD), MH763749^∗^, MH822264^∗^, MH752677^∗^, MH796626^∗^. ***Witheringia macrantha*** (Standl. & C.V. Morton) Hunz., COSTA RICA, Monteverde, *Bohs 2512* (UT), AY665857, MH822265^∗^, EU581071, MH796627^∗^. ***Witheringia meiantha*** (Donn. Sm.) Hunz., COSTA RICA, *Bohs 3015* (UT), AY665856, MH822266^∗^, EU581072, MH796628^∗^. ***Witheringia mexicana*** (B.L. Rob.) Hunz., NA, *Bohs 3294* (UT), MH763750^∗^, MH822267^∗^, na, MH796629^∗^. ***Witheringia mortonii*** Hunz., COSTA RICA, *Bohs 2594* (UT), MH763751^∗^, MH822268^∗^, MH752678^∗^, MH796630^∗^. ***Witheringia solanacea*** L’Hér., COSTA RICA, *Bohs 2416* (UT), na, na, EU581074, na. PANAMA, *D’Arcy 16399* (MO), DQ314164, DQ301517, na, DQ309474. ***Witheringia stellata*** (Greenm.) Hunz., MEXICO, *Stone 1522* (UT), MH763752^∗^, MH822269^∗^, MH752679^∗^, MH796631^∗^. ***Witheringia wurdackiana*** Benítez, VENEZUELA, Táchira, Fernández Feo, *Benítez de Rojas & Rojas 5433* (MO), MH763753^∗^, na, na, na.

## Appendices 2-10

uploaded in separate files.

## LITERATURE CITED

Aberer, A.J., D. Krompass, and A. Stamatakis. 2013. Pruning Rogue Taxa Improves Phylogenetic Accuracy: An Efficient Algorithm and Webservice. Systematic Biology 62: 162–166. Available at: http://www.ncbi.nlm.nih.gov/pmc/articles/PMC3526802/.

Averett, J.E. 1973. Biosystematic study of *Chamaesaracha* (Solanaceae). Rhodora 75: 325–365. Available at: http://www.jstor.org/stable/23311250.

Avino, M., E.M. Kramer, K. Donohue, A.J. Hammel, and J.C. Hall. 2012. Understanding the basis of a novel fruit type in Brassicaceae: conservation and deviation in expression patterns of six genes. EvoDevo 3: 20.

Beaulieu, J.M., and M.J. Donoghue. 2013. Fruit evolution and diversification in campanulid angiosperms. Evolution 67: 3132–3144.

Beaulieu, J.M., and B.C. O’Meara. 2016. Detecting hidden diversification shifts in models of trait-dependent speciation and extinction. Systematic Biology 65: 583–601.

Beckman, N.G., and H.C. Muller-Landau. 2011. Linking fruit traits to variation in predispersal vertebrate seed predation, insect seed predation, and pathogen attack. Ecology 92: 2131–2140.

Bolmgren, K., and O. Eriksson. 2005. Fleshy fruits–origins, niche shifts, and diversification. Oikos 109: 255–272.

Bouckaert, R., J. Heled, D. Kühnert, T. Vaughan, C.-H. Wu, D. Xie, M.A. Suchard, ET AL. 2014. BEAST 2: a software platform for Bayesian evolutionary analysis. PLoS Computational Biolology 10: e1003537.

Bremer, B., K. Andreasen, and D. Olsson. 1995. Subfamilial and tribal relationships in the Rubiaceae based on *rbcL* sequence data. Annals of the Missouri Botanical Garden 82: 383–397.

Burnham, K.P., and D.R. Anderson. 2002. Model selection and multimodel interference. Springer Verlag, New York, USA.

Cedeño, M.M., and D.M. Montenegro. 2004. Plan exportador, logistico y de comercializacion de uchuva al mercado de Estados Unidos para frutexpo SCI Ltda. Bachelor’s thesis, Facultad de Ingeniería, Pontificia Universidad Javeriana, Bogotá, Cundinamarca, Colombia.

Cueva Manchego, M.A., S.D. Smith, and S. Leiva González. 2015. A new and endangered species of *Iochroma* (Solanaceae) from the cloud forests of central Peru and its Phylogenetic position in Iochrominae. Phytotaxa 227: 147–157.

Darriba, D., G.L. Taboada, R. Doallo, and D. Posada. 2012. jModelTest 2: more models, new heuristics and parallel computing. Nature Methods 9: 772.

Davis, C.C., W.R. Anderson, and M.J. Donoghue. 2001. Phylogeny of Malpighiaceae: evidence from chloroplast ndhF and trnLF nucleotide sequences. American Journal of Botany 88: 1830–1846.

De-Silva, D.L., L.L. Mota, N. Chazot, R. Mallarino, K.L. Silva-Brandão, L.M.G. Piñerez, A.V.L. Freitas, ET AL. 2017. North Andean origin and diversification of the largest Ithomiine butterfly genus. Scientific reports 7: 45966.

Deanna, R., G.E. Barboza, and C. Carrizo García. 2017. Phylogenetic relationships of *Deprea*: New insights into the evolutionary history of physaloid groups. Molecular Phylogenetics and Evolution 119: 71–80. Available at: https://doi.org/10.10167j.ympev.2017.11.001.

Deanna, R., A. Orejuela, and G.E. Barboza. 2018. An updated phylogeny of *Deprea* (Solanaceae) with a new species from Colombia: interspecific relationships, conservation assessment and a key for Colombian species. Systematics and Biodiversity In press.

Doyle, J.J., and J.L. Doyle. 1987. A rapid procedure for DNA purification from small quantities of fresh leaf tissue. Phytochemical Bulletin 19: 11–15.

Drummond, A.J., S.Y.W. Ho, M.J. Phillips, and A. Rambaut. 2006. Relaxed phylogenetics and dating with confidence. PLoS Biology 4: e88.

Drummond, A.J., M. Kearse, J. Heled, R. Moir, T. Thierer, B. Ashton, A. Wilson, and S. Stones-Havas. 2006. Geneious v4.6.1. Biomatters, lAuckland.

Edgar, R.C. 2004. MUSCLE: multiple sequence alignment with high accuracy and high throughput. Nucleic Acids Research 32: 1792–1797.

Eriksson, O., E.M. Friis, and P. Löfgren. 2000. Seed size, fruit size, and dispersal systems in angiosperms from the Early Cretaceous to the Late Tertiary. The American Naturalist 156: 47–58.

Fernandez-Hilario, R., and S.D. Smith. 2017. A new species of *Saracha* (Solanaceae) from the Central Andes of Peru. PhytoKeys 85: 31–43.

FitzJohn, R.G. 2012. Diversitree: comparative phylogenetic analyses of diversification in R. Methods in Ecology and Evolution 3: 1084–1092.

Francis, J.K. 2000. *Hernandia sonora* L. Mago, toporite Hernandiaceae Familia de las hernandias. General Technical Report IITF 15: 260.

Fritz, S.A., and A. Purvis. 2010. Selectivity in Mammalian Extinction Risk and Threat Types: a New Measure of Phylogenetic Signal Strength in Binary Traits. Conservation Biology 24: 1042–1051. Available at: https://doi.org/10.1111/j.1523-1739.2010.01455.x.

Gautier-Hion, A., J.-M. Duplantier, R. Quris, F. Feer, C. Sourd, J.-P. Decoux, G. Dubost, ET AL. 1985. Fruit characters as a basis of fruit choice and seed dispersal in a tropical forest vertebrate community. Oecologia 65: 324–337.

Gernhard, T. 2008. The conditioned reconstructed process. Journal of Theoretical Biology 253: 769–778.

Givnish, T.J., J.C. Pires, S.W. Graham, M.A. McPherson, L.M. Prince, T.B. Patterson, H.S. Rai, ET AL. 2005. Repeated evolution of net venation and fleshy fruits among monocots in shaded habitats confirms a priori predictions: evidence from an *ndhF* phylogeny. Proceedings of the Royal Society of London B: Biological Sciences 272: 1481–1490.

Gottschling, M., and J.S. Miller. 2006. Clarification of the taxonomic position of *Auxemma, Patagonula*, and *Saccellium* (Cordiaceae, Boraginales). Systematic Botany 31: 361–367.

Hall, J.C., T.E. Tisdale, K. Donohue, A. Wheeler, M.A. AlYahya, and E.M. Kramer. 2011. Convergent evolution of a complex fruit structure in the tribe Brassiceae (Brassicaceae). American Journal of Botany 98: 1989–2003.

He, C., T. Münster, and H. Saedler. 2004. On the origin of floral morphological novelties. FEBSLetters 567: 147–151.

He, C., and H. Saedler. 2005. Heterotopic expression of*MPF2* is the key to the evolution of the Chinese lantern of *Physalis*, a morphological novelty in Solanaceae. Proceedings of the National Academy of Sciences 102: 5779–5784.

He, C., and H. Saedler. 2007. Hormonal control of the inflated calyx syndrome, a morphological novelty, in *Physalis*. The Plant Journal 49: 935–946.

Hepper, N.F. 1991. Old World *Withania* (Solanaceae): A taxonomic review and key to the species. *In* J. G. Hawkes, R. N. Lester, M. Nee, and N. Estrada [eds.], Solanaceae III: Taxonomy, Chemistry, Evolution, 211–227. Royal Botanic Gardens & Linnean Society of London, London, UK.

Hu, J.-Y., and H. Saedler. 2007. Evolution of the inflated calyx syndrome in Solanaceae. Molecular Biology and Evolution 24: 2443–2453.

Hua, X., and L. Bromham. 2016. Phylometrics: an R package for detecting macroevolutionary patterns, using phylogenetic metrics and backward tree simulation. Methods in Ecology and Evolution 7: 806–810.

Huelsenbeck, J.P., R. Nielsen, and J.P. Bollback. 2003. Stochastic mapping of morphological characters. Systematic Biology 52: 131–158.

Hunziker, A. 1969. Estudios sobre Solanaceae V. Contribución al conocimiento de *Capsicum* y géneros afines (*Witheringia, Acnistus, Athenaea*, etc.). Primera parte. Kurtziana 5: 101–179.

Hunziker, A.T. 2001. Genera Solanacearum. A. R. G. Gantner Verlag, K.-G, Ruggell, Germany.

Iqbal, M., and A.K. Datta. 2007. Cytogenetic studies in *Withania somnífera* (l.) Dun. (Solanaceae). Cytologia 72: 43–47.

Khan, M.R., J.-Y. Hu, S. Riss, C. He, and H. Saedler. 2009. MPF2-like-A MADS-box genes control the inflated calyx syndrome in *Withania* (Solanaceae): roles of Darwinian selection. Molecular Biology and Evolution 26: 2463–2473.

Khan, M.R., J. Hu, and G.M. Ali. 2012a. Reciprocal loss of CArG-boxes and auxin response elements drives expression divergence of*MPF2*-Like MADS-box genes controlling calyx inflation. PLoS One 7: e42781.

Khan, M.R., J. Hu, and C. He. 2012b. Plant hormones including ethylene are recruited in calyx inflation in Solanaceous plants. Journal of Plant Physiology 169: 940–948.

Khan, M.R., I.U. Khan, and G.M. Ali. 2013. *MPF2*-Like MADS-Box Genes Affecting Expression of SOC1 and MAF1 are Recruited to Control Flowering Time. Molecular Biotechnology 54: 25–36. Available at: https://doi.org/10.1007/s12033-012-9540-9.

Knapp, S. 2002. Tobacco to tomatoes: a phylogenetic perspective on fruit diversity in the Solanaceae. Journal of Experimental Botany 53: 2001–2022.

Lagomarsino, L.P., F.L. Condamine, A. Antonelli, A. Mulch, and C.C. Davis. 2016. The abiotic and biotic drivers of rapid diversification in Andean bellflowers (Campanulaceae). New Phytologist 210: 1430–1442.

LarsonJohnson, K. 2016. Phylogenetic investigation of the complex evolutionary history of dispersal mode and diversification rates across living and fossil Fagales. New Phytologist 209: 418–435.

Leiva González, S., R. Deanna, and J.J. Gavilán. 2013. Tres nuevas especies de *Iochroma* Bentham (Solanaceae) del Norte del Perú. Arnaldoa 20: 25–44.

Leiva Gonzalez, S., P. Lezama Asencio, and V. Quipuscoa Silvestre. 2003. *Iochroma salpoanum* y *I. squamosum* (Solanaceae: Solaneae) dos nuevas especies andinas del norte del Perú. Arnaldoa 10: 95–104.

Leiva González, S., and P. Lezama. 2005. *Iochroma albianthum* e *Iochroma ayabacense* (Solanaceae: Solaneae) dos nuevas especies del Departamento de Piura, Perú. Arnaldoa 12: 72–80.

Lezama Escobedo, K., E. Pereyra Villanueva, S. Limo Cruz, and S. Leiva Gonzalez. 2007. *Iochroma smithianum* (Solanaceae) una nueva especie del Departamento La Libertad, Peru. Arnaldoa 14: 23–28.

Li, H.-Q., P. Gui, S.-Z. Xiong, and J.E. Averett. 2013. The generic position of two species of tribe Physaleae (Solanaceae) inferred from three DNA sequences: A case study on *Physaliastrum* and *Archiphysalis*. Biochemical Systematics and Ecology 50: 82–89.

Lomáscolo, S.B., D.J. Levey, R.T. Kimball, B.M. Bolker, and H.T. Alborn. 2010. Dispersers shape fruit diversity in *Ficus* (Moraceae). Proceedings of the National Academy of Sciences 107: 14668–14672.

Marcussen, T., and A.S. Meseguer. 2017. Species-level phylogeny, fruit evolution and diversification history of *Geranium* (Geraniaceae). Molecular Phylogenetics and Evolution 110: 134–149. Available at: http://www.sciencedirect.com/science/article/pii/S1055790317302130.

Mason-Gamer, R.J., and E.A. Kellogg. 1996. Testing for phylogenetic conflict among molecular data sets in the tribe Triticeae (Gramineae). Systematic Biology 45: 524–545.

Miller, M.A., W. Pfeiffer, and T. Schwartz. 2010. Creating the CIPRES Science Gateway for inference of large phylogenetic trees. *In* Gateway Computing Environments Workshop (GCE), 1–8, Ieee.

Ng, J., and S.D. Smith. 2014. How traits shape trees: new approaches for detecting character statedependent lineage diversification. Journal of Evolutionary Biology 27: 2035–2045.

Olmstead, R.G., L. Bohs, H. Abdel Migid, E. Santiago-Valentín, V.F. Garcia, and S.M. Collier. 2008. A molecular phylogeny of the Solanaceae. Taxon 57: 1159–1181.

Ortiz-Ramírez, C.I., S. Plata-Arboleda, and N. Pabón-Mora. 2018. Evolution of genes associated with gynoecium patterning and fruit development in Solanaceae. Annals of Botany 121: 1211–1230. Available at: http://dx.doi.org/10.1093/aob/mcy007.

Pabón-Mora, N., G.K.-S. Wong, and B.A. Ambrose. 2014. Evolution of fruit development genes in flowering plants. Frontiers in Plant Science 5: 300.

Padmaja, H., S. Sruthi, and M. Vangalapati. 2014. Review on *Hibiscus sabdariffa*-A valuable herb. International Journal of Pharmacy & Life Sciences 5: 3747–3752.

Paradis, E., J. Claude, and K. Strimmer. 2004. APE: Analyses of Phylogenetics and Evolution in R language. Bioinformatics 20: 289–90.

Paton, A. 1990. A global taxonomic investigation of *Scutellaria* (Labiatae). Kew Bulletin 45: 399–450.

Poczai, P., and J. Hyvönen. 2011. Identification and characterization of plastid *trnF* (GAA) pseudogenes in four species of *Solanum* (Solanaceae). Biotechnology letters 33: 2317.

Posada, D., and K.A. Crandall. 1998. Modeltest: testing the model of DNA substitution. Bioinformatics 14: 817–818.

Preston, J.C., and L.C. Hileman. 2009. Developmental genetics of floral symmetry evolution. Trends in Plant Science 14: 147–154.

Rambaut, A. 2016. FigTree, version 1.4.3. Computer program and documentation distributed by the author, website: http://tree.bio.ed.ac.uk/software/figtree/ [accessed 20 June 2017].

Rambaut, A., A.J. Drummond, D. Xie, G. Baele, and M.A. Suchard. 2018. Posterior summarisation in Bayesian phylogenetics using Tracer 1.7. Systematic Biology 67: 901–904.

Revell, L.J. 2012. phytools: an R package for phylogenetic comparative biology (and other things). Methods in Ecology and Evolution 3: 217–223.

Riss, S. 2009. Isolation and analysis of *MPF2*-like MADS-box genes from Physaleae and characterization of their cis-regulatory regions. Ph.D. dissertation, Universität zu Köln, Köln, Germany.

Särkinen, T., L. Bohs, R.G. Olmstead, and S. Knapp. 2013. A phylogenetic framework for evolutionary study of the nightshades (Solanaceae): a dated 1000-tip tree. BMC Evolutionary Biology 13: 214–229.

Särkinen, T., S. Kottner, W. Stuppy, F. Ahmed, and S. Knapp. 2018. A new commelinid monocot seed fossil from the early Eocene previously identified as Solanaceae. American Journal of Botany 105: 95–107.

Sawyer, N.W. 2001. New species and combinations in *Larnax* (Solanaceae). Novon 11: 460–471.

Schliep, K.P. 2011. phangorn: phylogenetic analysis in R. Bioinformatics 27: 592–593. Available at: http://dx.doi.org/10.1093/bioinformatics/btq706.

Shaw, J. 2018a. *Iochroma* reshuffle. The Plantsman 17: 124–125.

Shaw, J. 2018b. Response from Julian Shaw, Senior Registrar, RHS Botany Department. The Plantsman 17: 200.

Smith, S.D., C. Ane, and D.A. Baum. 2008. The role of pollinator shifts in the floral diversification of *Iochroma* (Solanaceae). Evolution 62: 793–806. Available at: http://www.ncbi.nlm.nih.gov/pubmed/18208567.

Smith, S.D., and D.A. Baum. 2006. Phylogenetics of the florally diverse andean clade Iochromidae (Solanaceae). American Journal of Botany 93: 1140–1153.

Sobel, J.M., and M.A. Streisfeld. 2013. Flower color as a model system for studies of plant evo-devo. Frontiers in Plant Science 4: 321.

Stamatakis, A. 2014. RAxML version 8: a tool for phylogenetic analysis and post-analysis of large phylogenies. Bioinformatics 30: 1312–1313.

Stenz, N.W.M., B. Larget, D.A. Baum, and C. Ané. 2015. Exploring tree-like and nontree-like patterns using genome sequences: an example using the inbreeding plant species *Arabidopsis thaliana* (L.) Heynh. Systematic Biology 64: 809–823.

Tamura, K., G. Stecher, D. Peterson, A. Filipski, and S. Kumar. 2013. MEGA6: molecular evolutionary genetics analysis version 6.0. Molecular Biology and Evolution 30: 2725–2729.

Tewksbury, J.J., and G.P. Nabhan. 2001. Seed dispersal: directed deterrence by capsaicin in chillies. Nature 412: 403.

Thiers, B. 2017. Index Herbariorum: A global directory of public herbaria and associated staff, [online] Website http://sweetgum.nybg.org/science/ih/. [accessed 6 June 2018].

Tiffney, B.H. 1984. Seed size, dispersal syndromes, and the rise of the angiosperms: evidence and hypothesis. Annals of the Missouri Botanical Garden 71: 551–576.

Traveset, A. 1998. Effect of seed passage through vertebrate frugivores’ guts on germination: a review. Perspectives in Plant Ecology, Evolution and Systematics 1: 151–190. Available at: http://www.sciencedirect.com/science/article/pii/S1433831904700104.

Turner, B.L. 2015. Taxonomy of *Chamaesaracha* (Solanaceae). Phytologia 97: 226–245.

Vaidya, G., D.J. Lohman, and R. Meier. 2011. SequenceMatrix: concatenation software for the fast assembly of multi gene datasets with character set and codon information. Cladistics 27: 171–180.

Vander Wall, S.B. 2001. The evolutionary ecology of nut dispersal. The Botanical Review 67: 74–117.

Wang, L., J. Li, J. Zhao, and C. He. 2015. Evolutionary developmental genetics of fruit morphological variation within the Solanaceae. Frontiers in Plant Science 6: 248.

Whitson, M., and P.S. Manos. 2005. Untangling *Physalis* (Solanaceae) from the Physaloids: A Two-Gene Phylogeny of the Physalinae. Systematic Botany 30: 216–230.

Wilf, P., M.R. Carvalho, M.A. Gandolfo, and N.R. Cúneo. 2017. Eocene lantern fruits from Gondwanan Patagonia and the early origins of Solanaceae. Science 355: 71–75.

Zamberlan, P.M., I. Rodrigues, G. Mäder, L. Castro, J.R. Stehmann, S.L. Bonatto, and L.B. Freitas. 2015. Reevaluation of the generic status of*Athenaea* and *Aureliana* (Withaniinae, Solanaceae) based on molecular phylogeny and morphology of the calyx. Botanical Journal of the Linnean Society 177: 322–334.

Zamora-Tavares, M. del P., M. Martínez, S. Magallón, L. Guzmán-Dávalos, and O. Vargas-Ponce. 2016. *Physalis* and physaloids: A recent and complex evolutionary history. Molecular Phylogenetics and Evolution 100: 41–50.

Zhang, J., M.R. khan, Y. Tian, Z. Li, S. Riss, and C. He. 2012. Divergences of *MPF2*-like MADS-domain proteins have an association with the evolution of the inflated calyx syndrome within Solanaceae. Planta 236: 1247–1260.

